# Structural insights and rational design of *Pseudomonas putida* KT2440 Omega transaminases for enhanced biotransformation of (*R*)-Phenylacetylcarbinol to (1*R*, 2*S*)-Norephedrine

**DOI:** 10.1101/2024.10.08.617172

**Authors:** Parijat Das, Santosh Noronha, Prasenjit Bhaumik

## Abstract

Omega transaminases (ω-TAs) can mediate the chiral amination of several unnatural substrates without the requirement of an α-COOH group, and are highly relevant in the production of several pharmaceutical intermediates of commercial interest. Development of better variants of ω-TAs are hence essential for their industrial uses. We have studied the active site architecture of the wild-type ω-TAs, to develop engineered enzymes for enhancing the biotransformation of (*R*)-Phenylacetylcarbinol to (1*R*, 2*S*)-Norephedrine. Two such ω-TAs (TA_5182 and TA_2799) from *P. putida* KT2440 strain were overexpressed and purified as recombinant proteins. Crystal structures of TA_5182 were solved in two conformations, and significant movements of two highly flexible loops were observed in these different states. The TA_2799 structure was determined in the co-factor bound state with a PLP molecule covalently bonded to the catalytic K286 as an internal aldimine. Enzyme assays indicated that TA_2799 required significantly higher concentrations of co-factor than TA_5182 to achieve satisfactory biotransformation of (*R*)-PAC. A key mutation of L322F in TA_2799 drastically reduced the co-factor dependency of the TA_2799_L322F mutant enzyme, and the mutant remained active for 96h at 30°C. The crystal structure of the mutant enzyme revealed an asparagine residue that mediates a hydrogen bonding network at the dimeric interface of the enzyme and is absent in TA_5182. The TA_5182_G119N mutant also showed enhanced co-factor affinity. The results of our studies will help generate *Pseudomonad* ω-TAs and ω-TAs from other organisms with high efficiency for asymmetric synthesis, to be used in host systems for optimal large-scale industrial biotransformation.

## INTRODUCTION

Transaminases (TA) or aminotransferases are a class of enzymes that catalyze the exchange of an amine (-NH_2_) group with the keto (C=O) group between an amino acid and an α-keto acid. These enzymes are highly stereo-selective and have two substrate binding pockets of varied sizes along with a cofactor pyridoxal 5ʹ-phosphate (PLP). During transamination, the binding and involvement of PLP molecule to form a Schiff’s base intermediate is one of the key steps in catalysis (1). The α- and β-TAs require a -COOH moiety for the catalytic reaction while ω-TAs can catalyze the transamination reaction without the requirement of the -COOH moiety. Therefore, ω-TAs can accept a wide range of donor and acceptor substrates leading to their potential use to synthesize commercially important compounds(2). Due to high enantiomeric selectivity and environmentally friendly concise reactions, transaminases have high industrial applications, especially in the pharmaceutical industry (3).

In recent years, biocatalytic routes have emerged as a promising approach to produce various bulk and fine chemicals, including pharmaceuticals (4). Among the significant advancements in this area, the biocatalytic synthesis of chiral amines has garnered considerable attention. Chiral amines, with their diverse applications as drugs and chiral auxiliaries in organic synthesis, play a crucial role in the pharmaceutical and chemical industries. One such industrially important chiral amine is (1*R,* 2*S*)-Norephedrine (1*R,* 2*S*)-NE). (1*R*, 2*S*)-NE and its derivatives are primarily used as chiral auxiliaries or chiral catalysts in various asymmetric reactions (5, 6). The synthesis of (1*R*, 2*S*)-NE from (*R*)-Phenylacetylcarbinol ((*R*)-PAC) *via* chemical routes pose several challenges, including low enantioselectivity and the requirement for multiple synthesis steps involving harsh reaction conditions (4, 5). As a result, researchers have turned to biocatalytic strategies, utilizing transaminases as stereoselective biocatalysts for chiral amine synthesis (6–8) (Figure 1). Although transaminases are important in several industrial applications, most reported transaminases suffer from insufficient operational or kinetic stability at high substrate concentrations, denaturing solvents, and elevated temperatures (9, 10). Hence, there is a need for the development of better variants of transaminases for industrial use. Research works have been focused on understanding the molecular basis of the kinetic instability of transaminases for their rationale engineering. Studies on the well-characterized transaminase from *C. violaceum* (*Cv-*TA) indicate that PLP binding, especially the binding of the 5ʹ-phosphate in the phosphate group binding cup (PGBC) acts as the driving factor for the structural rearrangements in the enzyme (11, 12). In this context, the affinity of transaminases towards the cofactor PLP plays a key role in maintaining the stability of the enzyme (9). In bacterial fold type I ω-TAs, two monomers are required to form the active site and the PLP binding pocket. The binding of the cofactor depends on the dynamics of the two flexible loops. The N-terminal loop acts as a “lid” covering the aromatic moiety of the PLP molecule, and it interacts with the C-terminal loop (termed as the “roof”) from the other monomer to accommodate the cofactor molecule in the active site of the enzyme. Increasing the affinity of the enzyme towards PLP enhances its stability and biophysical properties (9, 13–15). Over the last decade, studies have been focused on (*S*)-selective ω-TAs belonging to Fold type I from several organisms, like *C. violaceum* (11)*, V. fluvalis* (16)*, P. putida* (17), and *H. elongata* (18) that highlight some of the key residues that mediate polar interactions with the co-factor molecule and the movement of the loop during transition of the enzyme from its apo to holo form. However, the effect of the loop movement on enzyme activity and cofactor affinity is still not well understood. Transaminases have shown promise in efficiently consuming the substrate (*R*)-PAC for (1*R*, 2*S*)-NE synthesis. Among them, two transaminases TA_2799 (Uniprot ID: Q88J50) and TA_5182 (Uniprot ID: Q88CJ8) from *Pseudomonas putida* KT2440 have been previously reported for its catalytic potential towards formation of (1*R*, 2*S*)-NE (10, 17). However, there is a lack of detailed studies on the enzymatic synthesis of (1*R*, 2*S*)-NE using these specific enzymes. Therefore, to address this gap, comprehensive systematic studies of the transaminases from *Pseudomonas putida* KT2440 are essential to evaluate their suitability for the enzymatic synthesis of (1*R*, 2*S*)-NE.

**Figure 1:**
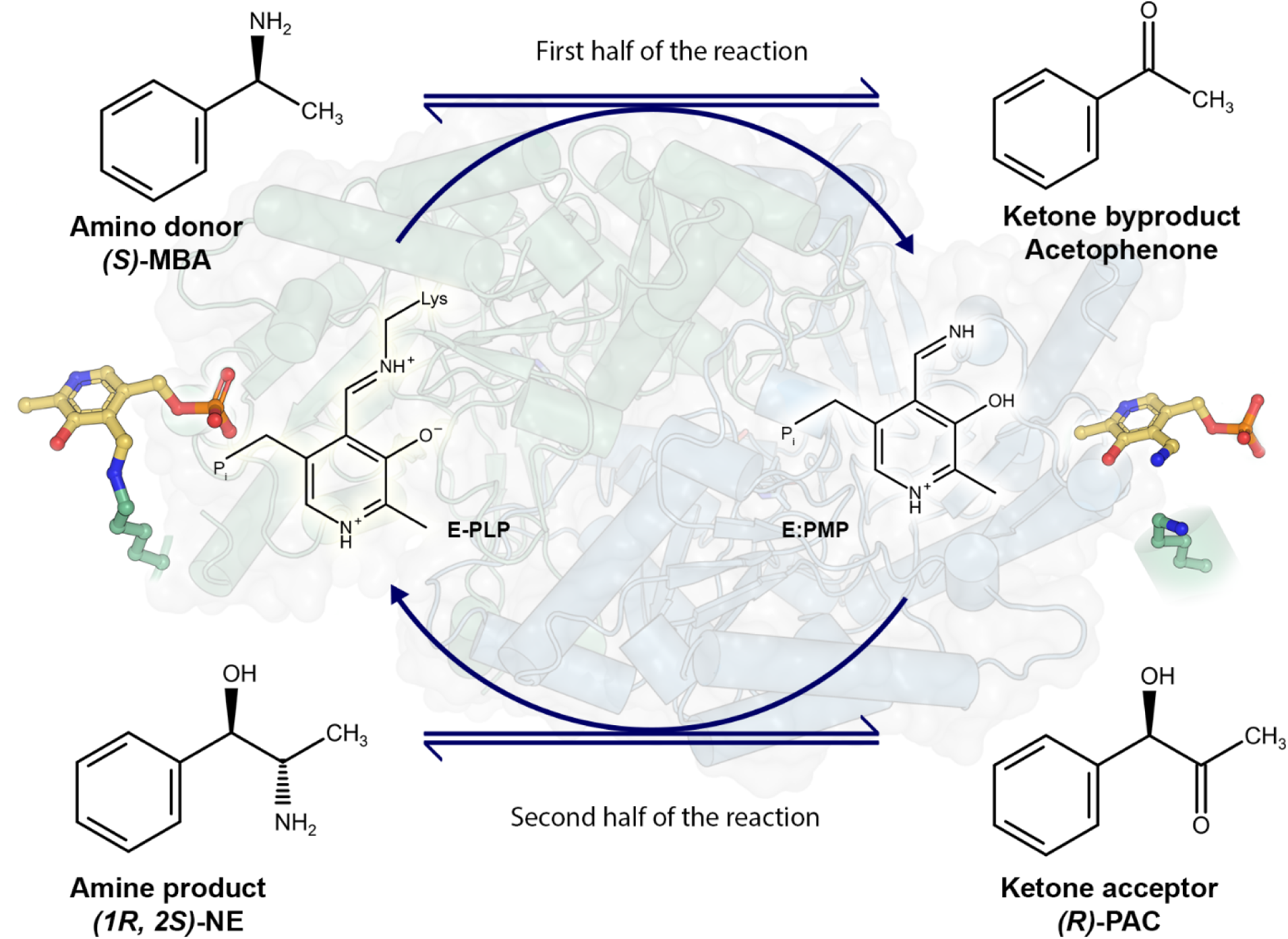
Reaction pathway of ω-TAs in the formation of (1*R*, 2*S*)*-*Norephedrine ((1*R*, 2*S*)*-*NE). In the first half of the reaction the amine group is transferred from the amine donor (*S*)-Methylbenzylamine ((*S*)-MBA) to the PLP molecule to form Pyridoxamine 5’-Phosphate (PMP) which is no longer covalently bound to the catalytic lysine of the enzyme. Acetophenone is produced as a by-product. In the second half of the reaction, the PMP molecule transfers the amine group to the acceptor (*R*)-PAC to form the amine product (1*R*, 2*S*)-NE and the enzyme is regenerated by forming the internal aldimine between PLP and the catalytic lysine.

In this work, we successfully overexpressed and characterized two recombinant transaminases namely TA_2799 and TA_5182 from *P. putida* KT2440 and solved their crystal structures. We determined the structure of TA_5182 in both relaxed (open) and closed states, mimicking the co-factor less and the co-factor bound form of the enzyme, respectively. The “roof” loop movement observed in these two forms revealed some important interactions essential for binding of the phosphate group of the cofactor molecule. Further, using structure based rational design approach key residues around the cofactor binding pocket of TA_2799 were mutated, and these single mutations exhibited increased affinity of the enzyme towards PLP. As a result, the mutants showed increased stability and enhanced biophysical properties, making them potential candidates for industrial bio-catalysis of NE. The results of this study would be applicable to other transaminases in general and aid in developing better ω-TAs for industrial uses.

## RESULTS

### Expression and purification of the wild-type TA enzymes and their stability

TA_2799 and TA_5182 were overexpressed in *E. coli* BL21 cells and purified using one-step Ni-NTA affinity chromatography and stored in a buffer containing 50 mM HEPES, 200 mM NaCl at pH 8.0 for biochemical characterization. TA_2799 posed a peculiar problem after purification of the enzyme. Even under incubation with 50 mM PLP at 4°C, the enzyme tended to precipitate within 24 h when concentrated over 5 mg/mL. Precipitation was hastened with increment in temperature and enzyme concentrations. On the other hand, TA_5182 remained stable in solution at high concentrations of 20 mg/mL. To address the problem of storage instability of TA_2799, we added glycerol to the freshly purified protein. It was found that a 12.5% (v/v) glycerol mixture effectively kept the enzyme stable in solution for at least 60 days, similar to the observation by Chen et al. with *C. violaceum* TA (13). This can be associated with the fact that glycerol prevents the aggregation of the enzyme molecules and thus prevents precipitation (19). We went ahead to crystallize the enzymes in their apo and PLP bound states to understand the structural basis of such a contrasting observation.

### Crystal structures of TA_2799 and TA_5182

The crystal structure of TA_2799 was determined at a resolution of 1.76 Å in the PLP-bound holo form. The structure of TA_5182 was solved in two different conformations in its apo state, at resolutions of 3.4 Å and 2.8 Å respectively. The crystal structure of L322F mutant of TA_2799 was solved at a resolution of 2.4 Å. The refinement statistics are summarized in Table 1.

**Table 1:**
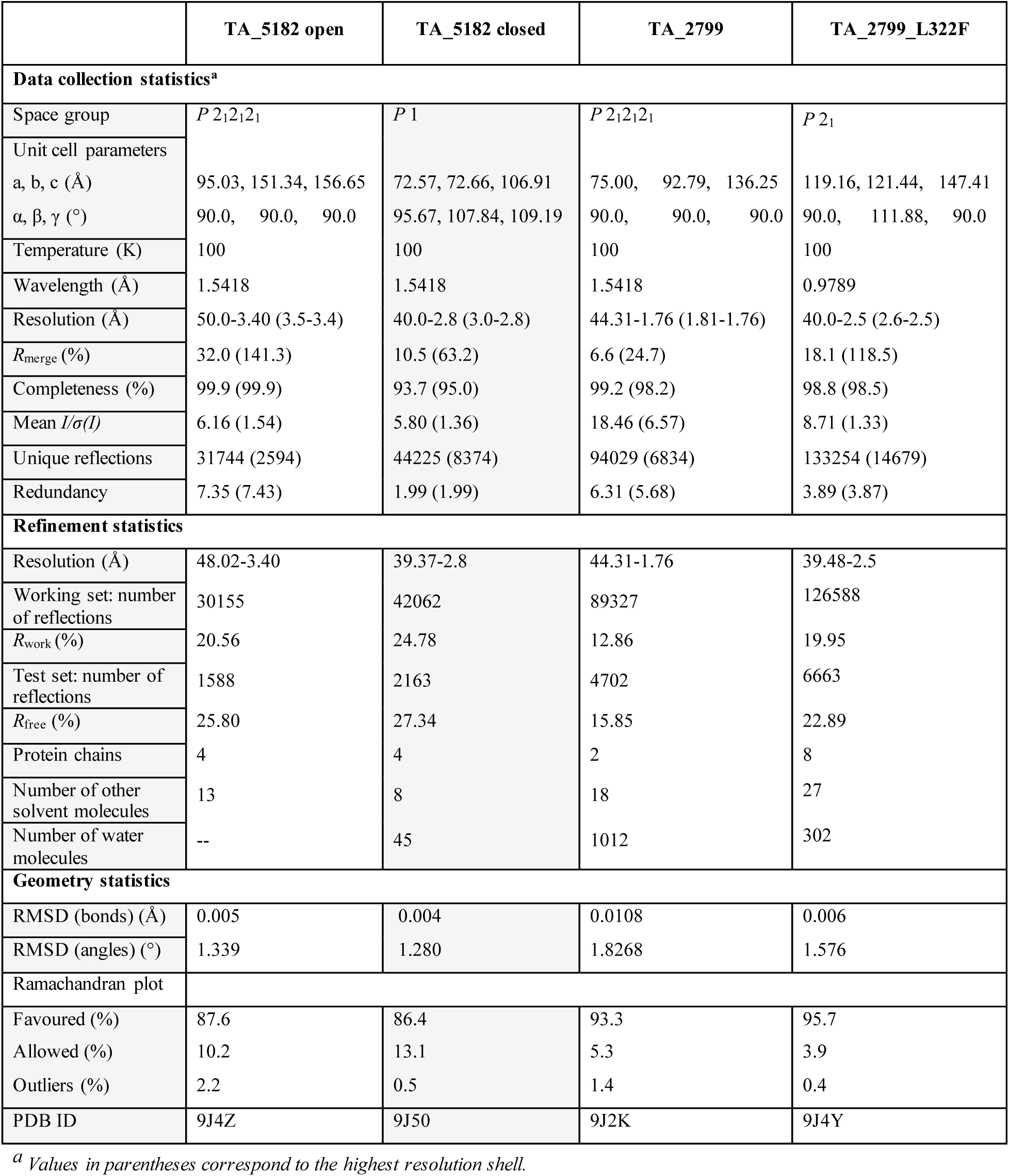
Data collection and refinement statistics of apo TA_5182 (open and closed forms), PLP bound TA_2799 and apo TA_2799 L322F mutant.

The overall structural fold of the TA_5182 and TA_2799 enzymes are similar to that observed for other class III Fold type I TAs with two α/β domains (Figure 2A). The enzymes function as a homodimer. For better understanding of the contributions of two subunits of the dimer, some of the residues are marked with a prime symbol to indicate the involvement of those residues from another monomer. Each monomer is composed of two major domains, the larger domain consisting of the residues 66-344, and the smaller N-terminal domain that consists of the residues 1-65 and the C-terminal region of 345-459 (Supplementary media 1). The large domain has an α/β/α sandwich topology, and is centered around a seven-stranded β-sheet as observed in other Fold type I ω-TAs (20). The small domain is made up of two β-sheet assemblies. The N-terminal β-sheet assembly is a three-stranded anti-parallel β-sheet instead of being four stranded as found in some ω-TAs like the *C. violaceum* TA and *P. aeruginosa* TAs where the last β-strand comes from the C-terminal part of the domain (21). This β-sheet assembly is capped by two flexible helices comprising of residues 1-32, electron densities of which were observed in the crystal structure of TA_2799 and the closed state of TA_5182. In the crystal structure of the open state of TA_5182, these residues could not be built due to poor electron density in this region, and hence were excluded from the final refined model. The C-terminal β-sheet is four-stranded with direction + - + + with a topology of +1, +2x, −1. As seen in other TAs, this sheet is shielded on one side from the solvent by three α-helices. The other side forms a crevice with the large domain which accommodates the enzyme’s active site.

**Figure 2:**
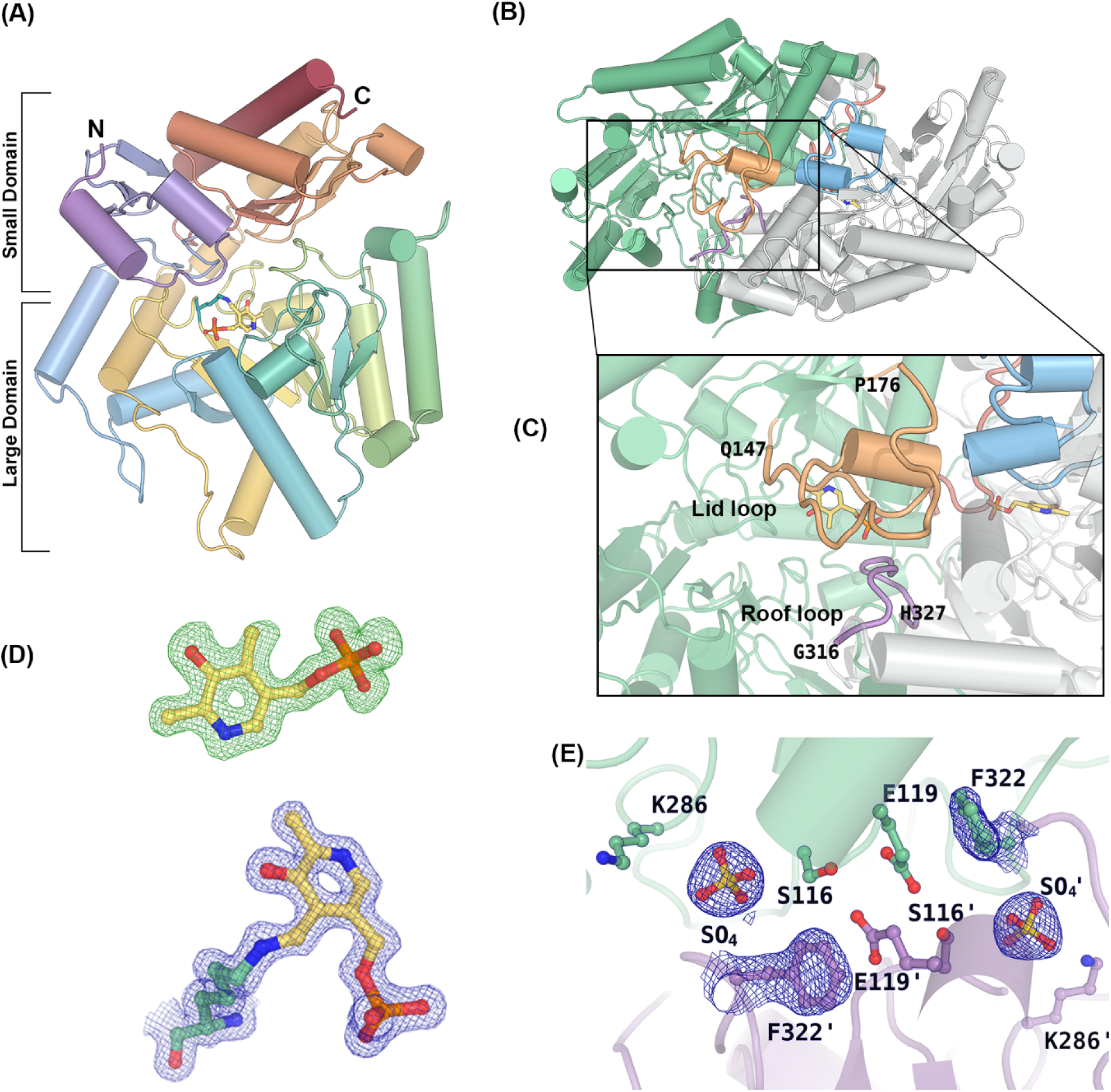
**(A)** Cartoon representation of the monomer of TA_2799 in rainbow spectrum showing the large and small domains. The PLP molecule covalently bound to the catalytic K286 is represented in ball and stick. **(B)** Dimeric view of TA_2799 showing the flexible ‘lid’ and ‘roof’ loops. **(C)** Zoomed in view of the lid (orange) and roof (purple) loops **(D)** σ-A weighted *F_o_-F_c_*omit map (green mesh) of PLP contoured at the 3.0 σ level superposed with the final refined model of PLP. Continuous electron density (blue mesh) can be observed after fitting the PLP covalently bound to the catalytic K286 in TA_2799 in the *2F_o_-F_c_* map contoured at 1.0 σ level. **(E)** Active site architecture of TA_2799_L322F mutant with sulphate ion at the phosphate group binding cup of the enzyme. The *2F_o_-F_c_* map contoured at 1.0 σ level (blue mesh) shows the electron density of the sulphate ion and F322 residue. The prime mark indicates the residues from another subunit of a dimer.

There are two highly flexible loops present in the structures of TA_2799 and TA_5182 (Figure 2B-C). These loops mediate the entry and exit of cofactor and substrate/byproduct into the active site of the enzyme, leading to significant conformational changes in the enzyme structure. We were able to crystallize TA_5182 in two different conformations which we term as the open and closed state (Figure 3A). On formation of the internal aldimine with the PLP molecule, the enzyme shifts from its open state to a more compact closed form. To date, there are very few studies where similar conformational differences have been reported, such as in the transaminase from *C. violaceum* (11) and *V. fluvialis* (16). In most of the apo form structures of the enzyme reported, the ‘roof’ region (residues 312-321 in TA_2799) that lines up the substrate entry channel is not visible due to high flexibility of these regions. In the apo structures of Fold type I transaminase enzymes from *Pseudomonas strains* (PDB ID: 5TI8, 6HX9), the roof structure is missing (17, 22). In our crystal structure of the apo state of TA_5182, the electron density of the roof loop is visible (Figure 3B). In the crystal structure of the open state of TA_5182 could not be visualized due to the inherent flexibility of this region. The ‘roof’ loop projects itself towards the PLP binding pocket on the binding of PLP in the active site of the enzyme. A sulphate ion can be observed at the PLP group phosphate binding pocket of the active site of this enzyme (Figure 3C). In the crystal structure of the closed state of TA_5182, electron density for the initial 32 residues and as well as that of the ‘roof’ and ‘lid’ loops are clearly visible. The conformation of the active site is similar to PLP-bound structures of ω-TAs; however, a phosphate ion can be seen at the phosphate group binding cup (PGBC) of the enzyme (Figure 3D).

**Figure 3:**
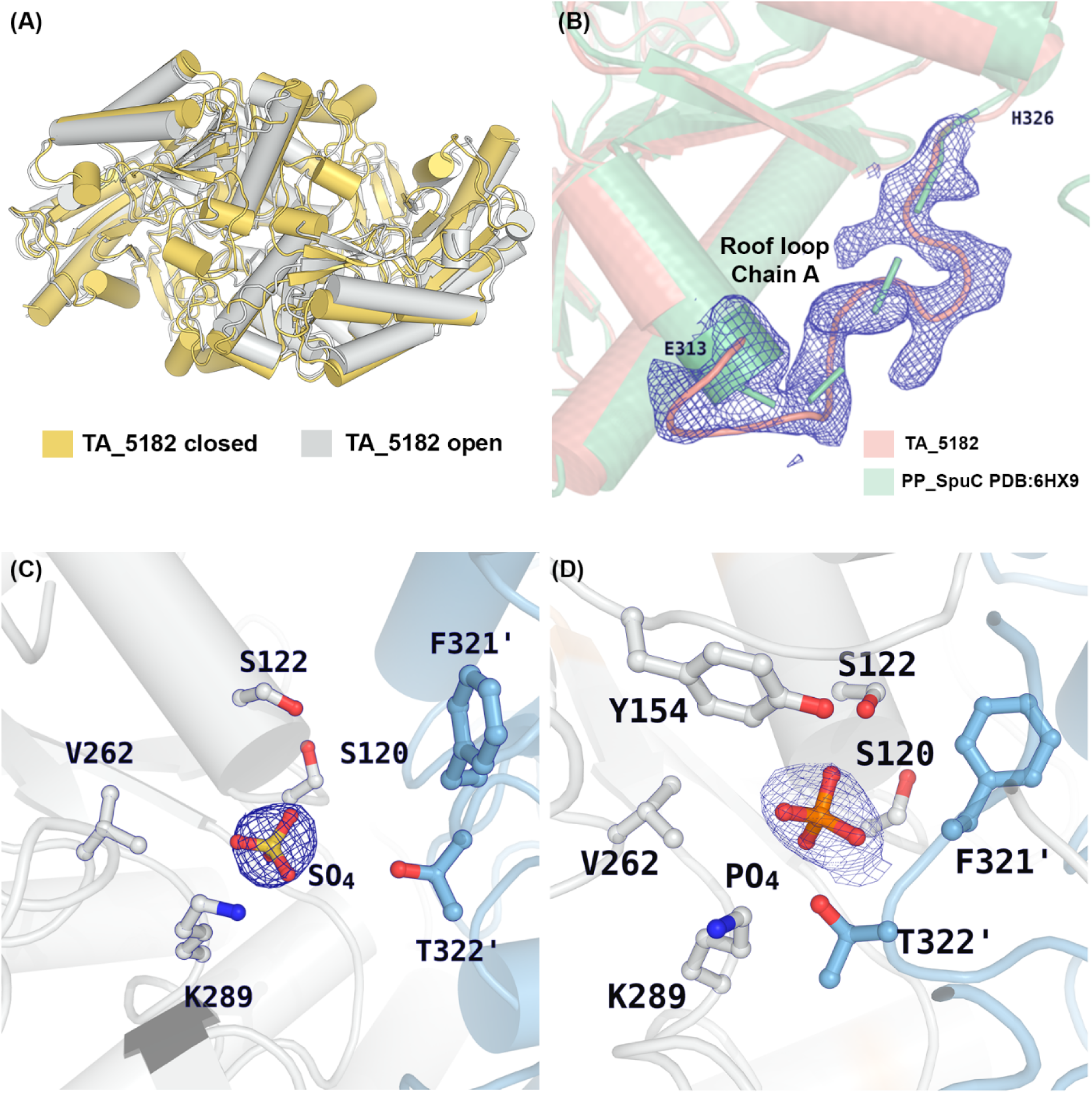
**(A)** Superimposition of the open (grey) and closed (yellow) crystal structures of dimeric TA_5182. **(B)** Superimposition of the open state of TA_5182 (salmon) and the putrescine TA from *P. putida* KT2440 (PDB ID: 6HX9) (green), showing the *2F_o_-F_c_* electron density map contoured at 1.0 σ level (blue mesh) of the roof loop in TA_5182. **(C)** Active site architecture of TA_5182 in the open state with sulphate ion electron density found at the phosphate group binding cup. **(D)** The active site architecture of the closed state of TA_5182 with phosphateion bound the phosphate group binding cup of the enzyme. The 2*F_o_-F_c_* electron density map of the phosphate and sulphate ions are represented in blue mesh (contour level: 1σ). The prime mark indicates the residues from another subunit of a dimer.

TA_2799 was crystallized in the PLP bound state. The conformation of the enzyme is almost identical to the closed state of TA_5182. Continuous high quality electron density could be seen for the covalently bound internal aldimine form of the PLP bound to the catalytic K286 residue (Figure 2D) in both the subunits. The TA_2799_L322F mutant was crystallized in its apo form, with a sulphate ion bound in the phosphate binding pocket of the enzyme (Figure 2E). These crystal structures and their analysis provide valuable information about the role of the different residues in and around the active site that mediate cofactor binding and the transamination reaction. More details about the roles of the active site residues and structural features of these ω-TAs in the catalytic reaction are discussed in the subsequent sections.

### Comparison of the active sites of TA_2799 and TA_5182

The high-resolution crystal structure of PLP bound TA_2799 provides insights into the architecture of the active site of this ω-TA. The active site is located in a deep cleft form by two monomers. Each dimer has two identical active sites which is formed by the contributions of the residues from both the subunits. The PLP molecule is linked to the ε-NH_2_ group of the catalytic K286 via a covalent Schiff base, and the pyrimidine ring of the cofactor molecule is sandwiched between the Y150 and I259 residues. The aromatic ring of the Y150 exhibits an edge-on interaction with the pyrimidine ring of the PLP molecule. The protonated pyrimidine nitrogen is stabilized by hydrogen bonded interactions with D257, and the hydroxyl group of the ring forms a water mediated hydrogen bond with E224. The phosphate group of PLP is stabilized by an intricate hydrogen bonding network, either directly or via water molecules, to S118, S285, Y150, and T323ʹ similar to the “phosphate group binding cup (PGBC)” described by Denesyuk et al (23).The superimposition of the apo structure of TA_5182 dimer in its closed state with that of the cofactor bound TA_2799 dimer shows that the overall fold of the enzymes is similar, having a root mean square deviation (RMSD) of 1.08 Å over 736 C-alpha atoms of the dimeric state of the enzymes. The enzymes TA_2799 and TA_5182 have a sequence identity of 38.49% (Supplementary figure 1), but the key residues that interact with the PLP molecule are identical except for I259 and L322 (Figure 4A and B). Comparison of TA_2799 with other known omega transaminases (9, 11, 14, 16, 17, 21, 24, 25) in literature shows the presence of a conserved valine at the I259 position of TA_2799 (Figure 4D). The presence of a slightly bulkier I259 may play a significant role in the occupancy of the external aldimine molecule in the active site pocket and might be the reason for the rapid conversion of PLP to PMP by the enzyme. On the other hand, the L322 position in TA_2799 is usually occupied by an aromatic amino acid (Figure 4D). In TA_5182, the position is occupied by F321, and the aromatic ring of phenylalanine contributes to a NH-π stacking interaction with an adjacent histidine residue, and an anion-π stacking with a conserved glutamate E123 on the other side which confers stability to the dimer interface (Figure 4C). Furthermore, in TA_2799, the glutamate residue interacts with an asparagine and a conserved serine residue to form an intricate hydrogen bonding network at the dimer interface. In TA_5182, the asparagine is replaced by a glycine residue (Figure 4E).

**Figure 4:**
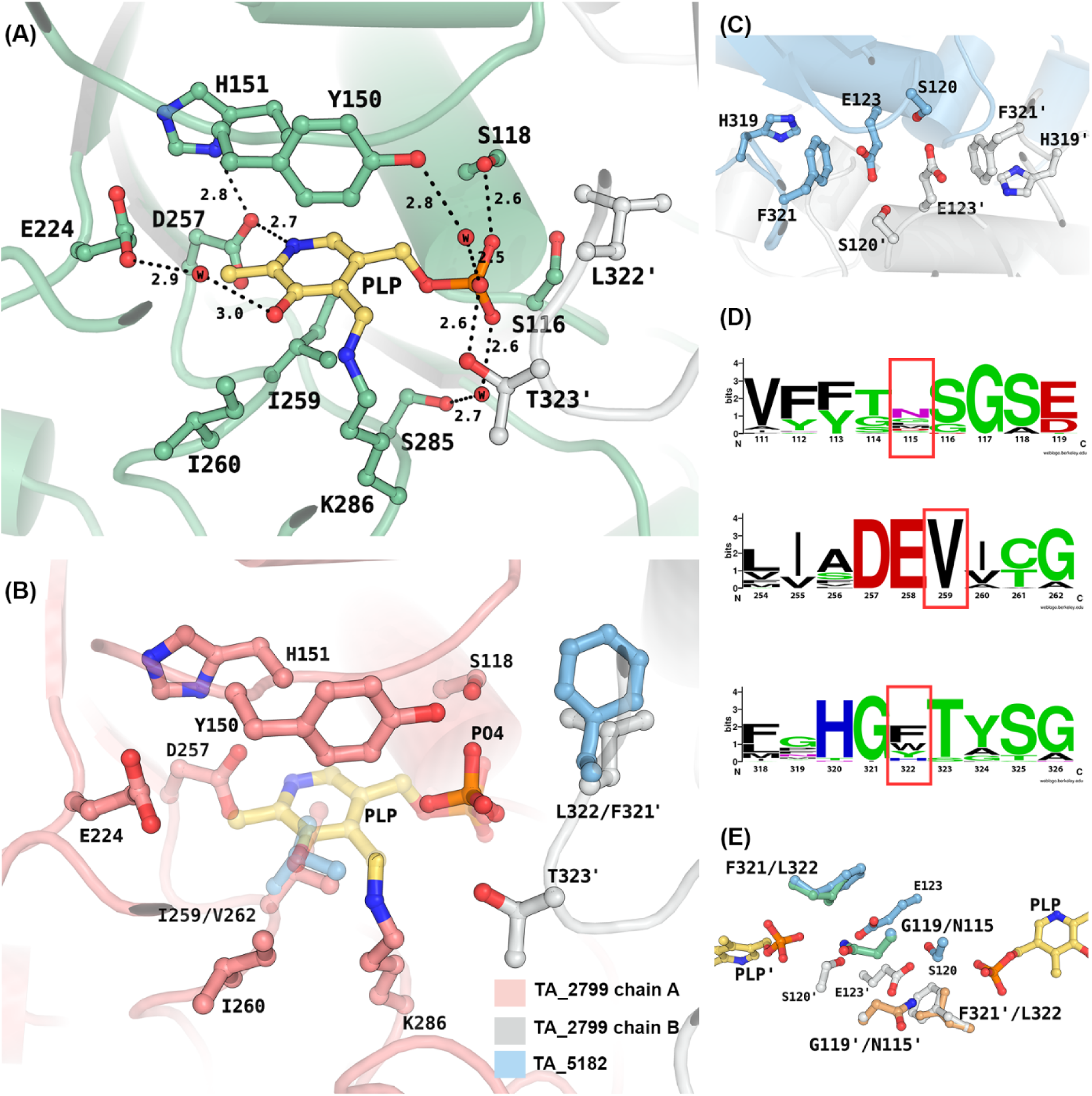
**(A)** Active site residues in PLP bound state of TA_2799 showing water mediated polar interactions between the residue sidechains and PLP. The PLP molecule is covalently bound to the catalytic K286. The black dotted lines represent the polar interaction distances in Å. **(B)** Superimposition of TA_2799 and the closed state of TA-5182 showing the differing residues (light blue). The chains from different monomers of TA_2799 are coloured in salmon and white. The PLP molecule present in TA_2799 is coloured in yellow, while the phosphate ion from TA_5182 is coloured in orange**(C)** Residues present at the dimeric interface of TA_5182 showing the E123 mediated anion-pi stacking interaction with F321. **(D)** Sequence logo representation of the conserved amino acids at the active site of bacterial ω-transaminases found in the PDB. The residues of interest are highlighted with red boxes. Residue numbers correspond to TA_2799 sequence. **(E)** Superimposition of TA_2799 and the closed state of TA_5182 showing the position of N115 and F321ʹ. The prime mark indicates the residues from another subunit of a dimer.

To assess the role of these residues in the cofactor binding and stability of omega transaminases, we constructed single mutants of TA_2799 and TA_5182. We compared the activity of the single mutants TA_2799_I259V, TA_2799_L322F and TA_5182_G119N with the wild-type enzymes to increase the stability and the affinity of the enzymes towards its cofactor PLP. Apart from this, we also evaluated the role of the Y323 in TA_5182, which is also somewhat conserved in ω-TAs (Figure 4D) by evaluating two mutants, TA_2799_L322F_C324Y and TA_5182_Y323C.

### Biochemical characterizations of the TA_2799, TA_5182 and their mutants

To evaluate the activity of the TA enzymes towards conversion of (*R*)-PAC to (1*R*, 2*S*)-NE, we used the Triphenyl Tetrazolium (TTC) based assay developed by Sehl et al. (8). For kinetic studies, pyruvate was used as the amine acceptor and (*S*)-MBA was used as the amine donor. TA_2799_I259V, surprisingly had very less activity (∼20%) compared to wild type TA_2799 (Supplementary figure 5).

The TA enzymes are known to demonstrate substrate inhibition (26). To understand the extent of substrate concentration that can be used for the TA enzymes for efficient catalysis, the enzymes were subjected to varying concentrations of the substrates keeping other parameters fixed. It was observed that TA_5182 has a better tolerance towards substrate inhibition in comparison to TA_2799. Interestingly, the ratio of the amino donor: amino acceptor needs to be at least 2:1 for efficient catalysis (>90% conversion) while using (*S*)-MBA as the amino donor (Figure 5A and B).

**Figure 5:**
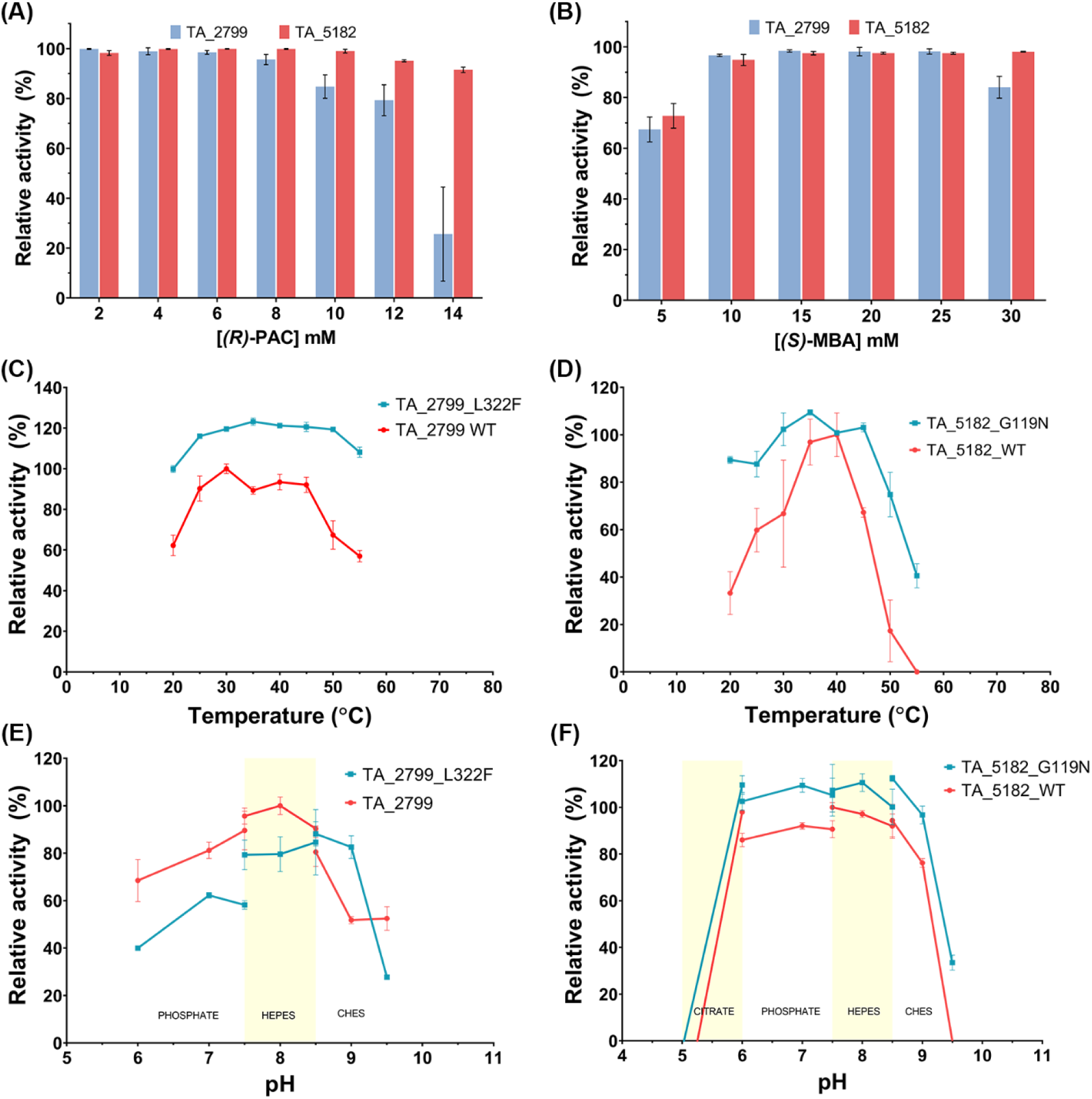
**(A)** Comparison of the conversion of varying (*R*)-PAC concentration by TA_5182 and TA_2799 in the presence of PLP with (*S*)-MBA as the amino donor**. (B)** Conversion of (*R*)-PAC by TA_5182 and TA_2799 in the presence of PLP with varying concentrationsof (*S*)*-*MBA. **(C)** Temperature optimum curve of TA_2799 and TA_2799_L322F **(D)** Temperature optimum curve of TA_5182 and TA_5182_G119N **(E)** pH optimum curve of TA_2799 and TA_2799_L322F **(F)** pH optimum curve of TA_5182 and TA_5182_G119N.

We studied the effect of temperature on the activities of the wild type and mutant enzymes. The TA_2799_L322F mutant showed a notable enhancement in enzymatic activity for the mutant variant across a range of temperatures compared to the wild-type enzyme (Figure 5C). The optimum activity for the wild-type TA_2799 enzyme was observed at 30°C, while the TA_2799_L322F mutant showed a similar activity across a wide range of temperature from 25-60°C. For TA_5182 and the TA_5182_G119N mutant, a similar temperature profile was observed (Figure 5D). The G119N mutant showed significantly higher activity at temperatures higher than 35°C as compared to the wild-type enzyme.

The pH dependent activities of the enzymes and their mutants were also measured. It was found that TA_2799 exhibits highest activity at a pH range of 7.5-8.5. The pH optima tend to shift towards alkaline pH in case of the L322F mutant (Figure 5E). TA_5182, on the other hand, was found to be active at the broader pH range. The optimum activity of the enzyme was found to be at pH 7.5, but it retained ∼80% of its activity at pH 6.0 as well. The activity dropped drastically to zero at pH 5.0 (Figure 5F). A similar trend was observed for the G119N mutant as well. More importantly, the mutant enzyme showed a higher relative activity compared to its wild-type counterpart.

### Cofactor release and PLP dependent conversion of (*R*)-PAC by ω-TAs

Loss of the aminated cofactor (PMP) formed during the first half of the reaction is a major cause for inactivation of transaminases (9, 26). A study by Schell et al. (27) showed that the PMP generated by the TA enzymes is released into the reaction medium due to the flexibility of the loops, as PMP is no longer covalently bound to the enzyme.

To investigate the cofactor release TA_2799 and TA_5182 were incubated with 1mM of PLP and 10mM of S-MBA and incubated at 30℃ in the dark in the presence and absence of 5mM amine acceptor (*R*)-PAC. PLP in solution has an Absorption maximum (Abs_Max_) at 408nm and PMP has an Abs_Max_ at 326nm. The spectra showed characteristic peaks at 408 and 326nm for PLP and PMP, respectively. The shorter roof region of TA_5182 was thought to be more flexible and thus caters to escape of the cofactor. However, when (*R*)-PAC was not present in the solution, it was seen that TA_2799 converted all the PLP into PMP, while TA_5182 still retained some of the PLP in the solution (Figure 6A and C). TA_2799 carries out this conversion much faster compared to TA_5182. In the cofactor binding pocket of TA_2799, however, as mentioned earlier, there are substitutions at the I259 in place of valine found in other ω-TAs. The presence of the bulkier isoleucine residue may play a significant role in release of the unbound external aldimine PMP from the cofactor binding pocket, thus leading to rapid conversion of PLP in the reaction mixture. However, the mutant TA_2799_L322F shows conversion of PLP to PMP at a slower rate than the wild-type enzyme, which indicates that F322 has a key role in retaining the PMP to the enzyme active site when the covalent bond has been broken (Figure 6B).

**Figure 6:**
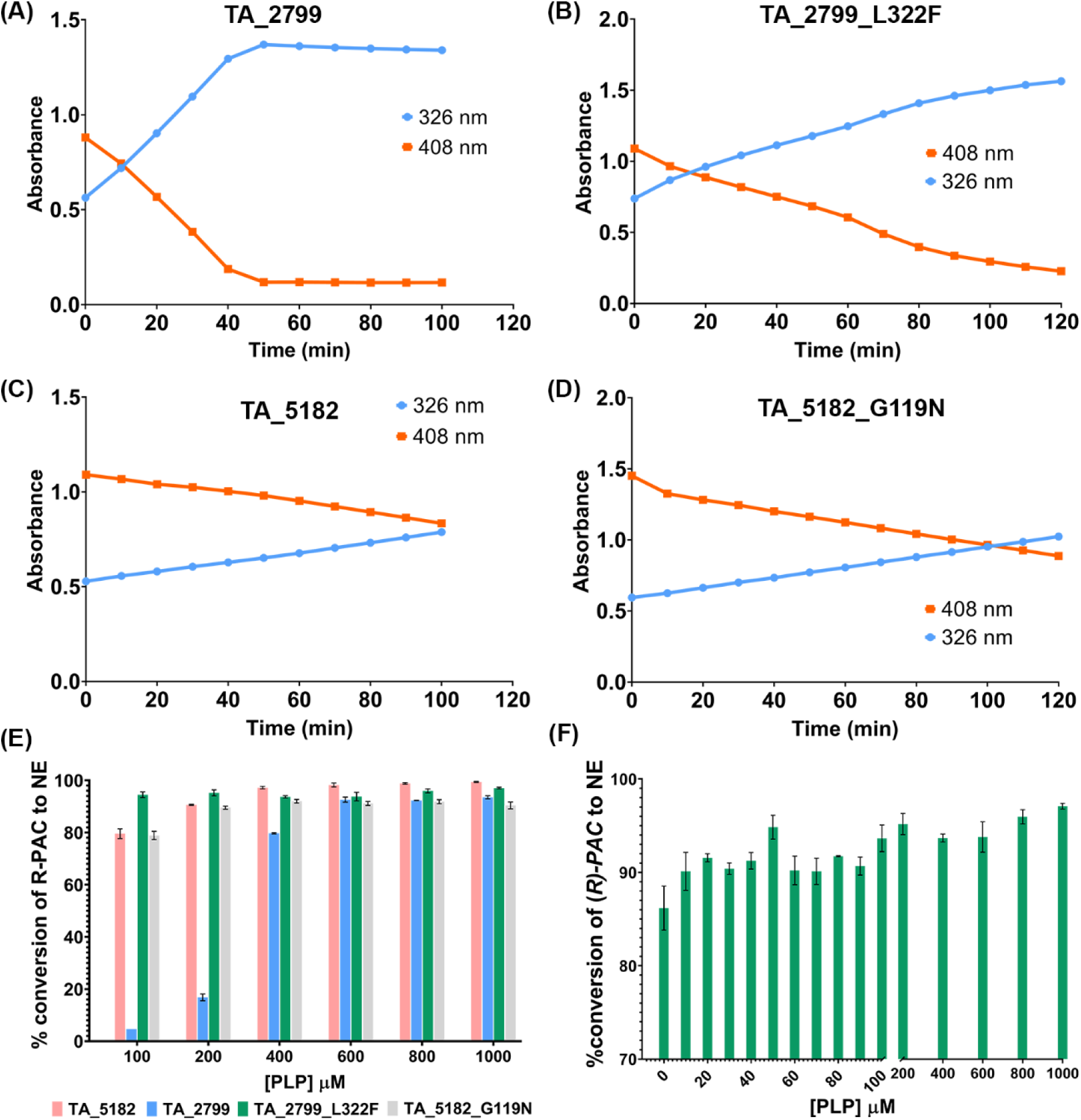
**(A)** Time dependent conversion of PLP to PMP by enzymes incubated with 1mM PLP and 10 mM *(S)*-MBA (amine donor) by TA_2799, **(B)** TA_2799_L322F, **(C)** TA_5182, and **(D)** TA_5182_G119N. Orange line represents the absorbance at 408 nm that denotes the amount of free PLP in solution, while the blue line denotes the absorbance of free PMP at 326nm. **(E)** Conversion of *(R)-*PAC in the presence of *(S)-*MBA by the TA enzymes at varying concentrations of PLP. **(F)** Conversion of *(R)-* PAC in the presence of *(S)*-MBA by TA_2799_L322F at varying concentrations of PLP.

The conversion of 5 mM (*R*)-PAC in the presence of 10 mM (*S*)-MBA at different concentrations of the cofactor PLP was also studied. TA_5182 showed efficient conversion at PLP concentrations of >200 μM, while TA_2799 required >400 μM of the cofactor in the reaction mixture for >80% conversion of R-PAC (Figure 6E). Since the TA_2799_I259V mutant showed decreased enzyme activity (Supplementary figure 5), the TA_2799_L322F mutant was probed for its PLP dependency. Interestingly, this mutant shows exceptionally good conversion efficiency, with >90% conversion of (*R*)-PAC even at PLP concentrations lower than 100 μM (Figure 6F).

### Thermal stability of TA_2799 and TA_5182 and their mutants with various additives

The wild type enzymes and TA_2799_L322F were assessed for the evaluation of their thermal stability with various additives added to the buffer. The Tm_app_ of wild type TA_5182 was found to be 57.5°C, while for wild type TA_2799 it was 56.6°C. Addition of glycerol and sucrose did not have a significant effect on the Tm_app_ of TA_5182, but for TA_2799, there was a significant increase in the Tm_app_ (Figure 7A and C). Sucrose has a better effect on increasing the stability of TA_2799 as compared to glycerol. In case of TA_5182, however, the increase in the amount of sucrose does not have any effect on the Tm_app_ of the enzyme. A similar trend was observed for the TA_5182_G119N mutant, where the Tm_app_ of the enzyme seems to be unaffected by the addition of sucrose and increased by 5°C on addition of 40% glycerol. A ∼5°C shift in the Tm_app_ of the mutant enzyme is observed over its wild-type counterpart (Figure 7D).

**Figure 7:**
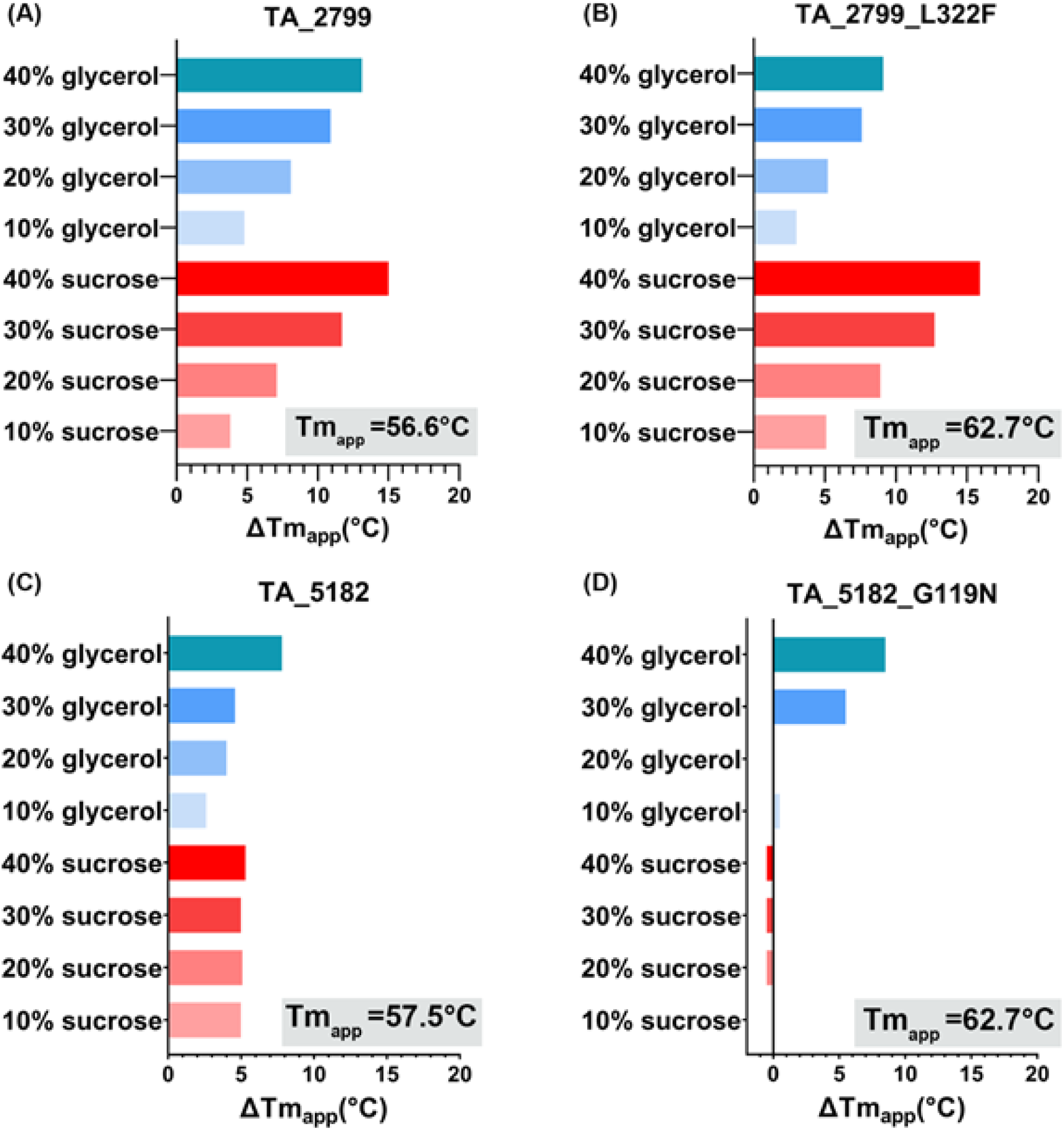
Shift in the Tm_app_ of **(A)** TA_2799, **(B)** TA_2799_L322F, **(C)** TA_5182 and **(D)** TA_5182_G119N in the presence of additives like glycerol and sucrose in varying concentrations. The Tm_app_ of the enzymes in the absence of any additives is denoted in the grey boxes.

The single mutant TA_2799_L322F also showed an Tm_app_ increase of 6.1°C over the wild-type enzyme (Figure 7B). Addition of glycerol and sucrose conferred a significant increase in the Tm_app_ of the mutant. The effect of sucrose on the Tm_app_ of the enzyme was more prominent, in the presence of 40% glycerol, the Tm_app_ increased by 9.1°C, while in the presence of 40% sucrose, the Tm_app_ increased by 15.1°C. The increased affinity of the enzyme to its cofactor PLP is the driving cause for the increase in the Tm of the mutant enzyme, as the binding of PLP has been shown to increase the stability of ω-TAs in several studies (9, 13–15). These results suggest that the mutant enzymes are more stable than the wild type, and the residues F322 (in TA_2799_L322F) and N119 (in TA_5182_G119N) play a key role in the stability of Fold type I ω-TAs.

### Storage stability of the transaminases

The transaminase enzymes were stored at 30°C without any external addition of PLP to measure the storage stability of the enzymes. TA_2799 lost its activity within 24h, while TA_5182 retained ∼40% its activity for 48h. The TA_5182_G119N mutant showed better tolerance compared its wild-type counterpart, as it retained 20% of its activity after 72h of storage. The TA_2799_L322F mutant showed the best increase in stability; it retained ∼70% of its activity even after 96h of storage (Figure 8). A possible explanation is that the mutant enzymes can capture the PLP cofactor from the cell lysate during the process of purification (as observed from the yellow colouration) which helps them maintain their tertiary structures. The binding of the PLP molecule at the PLP binding site reduces the conformational dynamics in the enzyme, as a result, the irreversible unfolding process is hindered.

**Figure 8:**
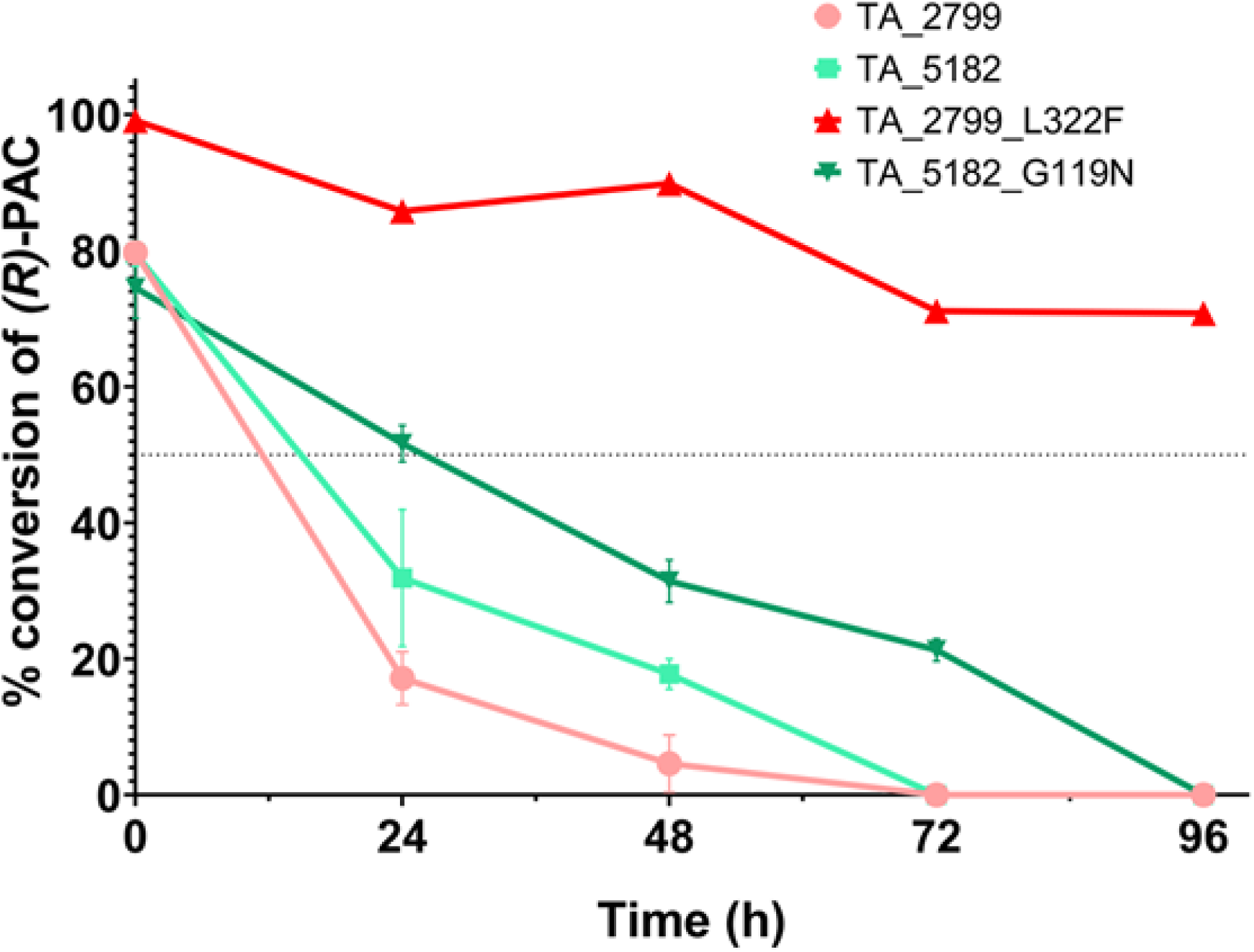
Conversion of *(R)*-PAC by TA_2799, TA_5182 and the mutant enzymes after storage at 30°C for different time periods. The mutant enzymes retain their activity better than their wild-type counterparts. The TA_2799_L322F mutant retains ∼70% of its activity even after 96 h of storage at 30°C.

### Kinetic parameters of the enzymes and their mutants

We evaluated the kinetic parameters of the TA enzymes and their mutants, as reported in Table 2. ω-TA enzymes are slow in their catalysis of unnatural substrates (25). It was seen that TA_2799 was ∼15 times faster than TA_5182 in terms of their rates of catalysis, although the affinity of TA_5182 for the amine donor substrate *(S)*-MBA was greater than that of TA_2799. The mutant TA_2799_L322F showed an increased affinity towards the amine donor, however, the V_max_ of the enzyme and the turnover number was reduced, probably because of the diminished movement of the dimeric interface. However, the catalytic efficiency of the mutant enzyme was calculated to be slightly greater than its wild-type counterpart. The TA_5182_G119N mutant also showed an increased affinity towards the amine donor (*S*)*-*MBA compared to the wild-type TA_5182, however, the turnover number of the mutant enzyme was reduced. However, the catalytic efficiency of the mutant enzyme was slightly enhanced due to the increased affinity towards the substrate.

**Table 2:** Kinetic parameters of the ω-TA enzymes against the amino donor (*S*)-MBA.

| Enzyme | $K_M$ (mM) | $V_{\max}$ (U mg<sup>-1</sup>) | $k_{\text{cat}}$ (min<sup>-1</sup>) | $k_{\text{cat}}/K_M$ (mM<sup>-1</sup> min<sup>-1</sup>) |
| --- | --- | --- | --- | --- |
| TA_2799 | 6.459 ± 0.621 | 0.104 ± 0.007 | 4.321 ± 0.279 | 0.668 |
| TA_2799_L322F | 1.068 ± 0.566 | 0.042 ± 0.001 | 1.777 ± 0.036 | 1.664 |
| TA_5182 | 1.089 ± 0.117 | 0.007 ± 0.0002 | 0.294 ± 0.007 | 0.269 |
| TA_5182_G119N | 0.347 ± 0.024 | 0.003 ± 0.0005 | 0.1468 ± 0.002 | 0.423 |

## DISCUSSION

Large scale industrial applications of the wild type transaminases are mostly constricted by operational instability and enzyme inactivation (9). Different approaches to engineer wild type transaminases have proven promising in increasing the yield of the desired amine product and improving the stability of the enzyme (28). The dependence of the enzyme on the co-factor PLP plays a key role in the applicability of these enzymes since external addition of the cofactor increases the cost of production (14). Moreover, cofactor binding enhances the stability and the catalytic activity of the enzymes (15). Our study hints at the possible factors that can lead to the development of ω-TAs with enhanced affinity towards the cofactor as well improved thermal stability of the engineered enzymes. Taking into account of the structural and biochemical data on TA_2799, TA_5182 and their variants, we have discussed the importance of some of the structural features that modulate the functional properties of ω-TAs.

### Loop movement and structural orientations of the F321 in TA_5182

The F321 (L322 in TA_2799) in TA_5182 is present in the highly flexible loop that is also known as the ‘roof’ of the PLP binding site (11). This loop projects itself towards the PLP binding pocket on the binding of PLP in the active site of the enzyme. Most of the apo-crystal structures of ω-TAs have been determined with a ligand which is either phosphate or a negatively charged ligand at the phosphate binding pocket of the active site of the enzyme. As described by Humble et al. in their study of ω-TA from *C. violaceum* (11), the position of the loop is vastly different in the one of the apo state of the enzyme (PDB ID: 4A6U), where the roof and lid loops show high flexibility. In the apo-open state of the enzyme, the F321 in TA_5182 initially is positioned near the surface of the enzyme and due to which the substrate entry channel of the enzyme is wider. On PLP binding, this residue moves by ∼14 Å orienting towards the α6 helix which has S120 and S122 from the other subunit (Figure 9A and B). This restructuring causes S120 and S122 to shift from their initial positions, and their interactions with a E123ʹ is broken, allowing S120 to coordinate the phosphate group of PLP (Figure 9C and supplementary media 2). The F321 itself gets stabilized by π-stacking interaction, sandwiched between a neighbouring H319 and the E123 (Figure 4C). In TA_2799, the side chain of L322 is not bulky enough to cause this change, and as a result the phosphate group of PLP may not be as tightly coordinated when the PLP molecule is bound to the enzyme. L322 cannot form the stacking interactions with H319 and G123 like aromatic amino acids. The effect of the L322F mutation is clearly observed in the thermal stability assay, as the mutant enzyme has an increased ∼6°C T_m_ over the wild-type TA_2799. Affinity of the enzyme towards the cofactor PLP is clearly enhanced, as observed by the yellow coloration of the mutant enzyme solution right after purification and the characteristic peak for PLP at 420nm seen in the absorbance spectra of the purified enzyme, similar to the report of Kwon et al. (29) (Supplementary figure 3A and 4A). The mutant enzyme shows efficient conversion of (*R*)-PAC to (1*R*, 2*S*)-NE even without external addition of any cofactor to the reaction mixture. We expected the presence of electron density of PLP in the crystal structure of the TA_2799_L322F mutant, however, only density of sulphate ion could be seen. It is possible that PLP was present at the active site of the enzyme, but not in the covalently bound internal aldimine state, and thus the sulphate ions from the crystallization mother liquor could compete and displace the PLP molecule out of the active site pocket.

**Figure 9:**
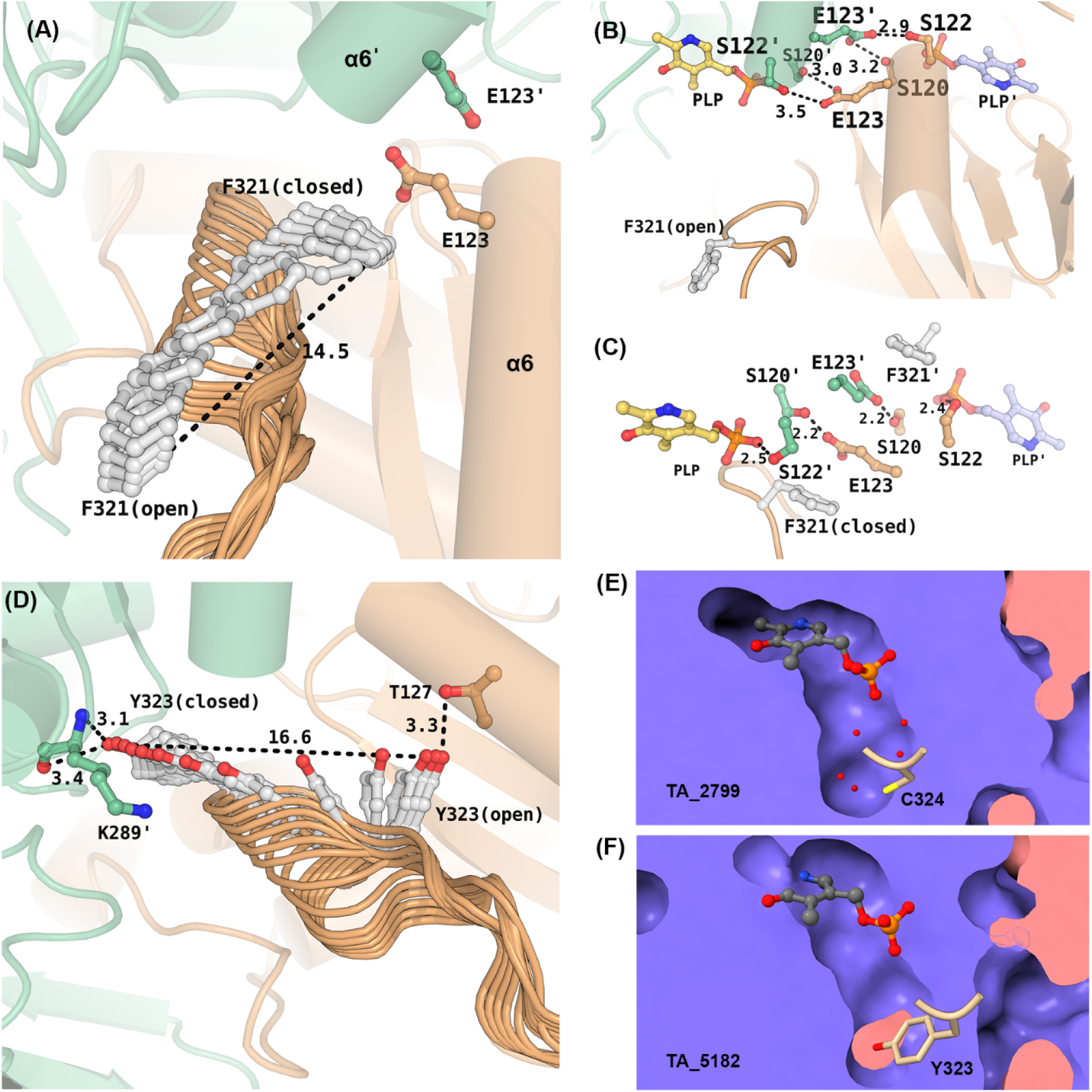
**(A)** Movement of F321 in TA_5182 from open to closed state. The E123 residue from each monomer is shown in ball and stick representation. The black dotted line represents the displacement of the F321 residue from open to closed state. **(B)** Interactions between E123 and S120 and S122 of the two subunits in the open state of TA_5182. **(C)** Interactions between E123 and S120 of the two subunits in the closed state of TA_5182. S122 is no longer in H-bonding distance with E123. The PLP molecule has been superposed from the crystal structure of TA_2799. **(D)** Movement of Y323 in TA_5182. Interactions between Y323 and T127 at initial open position. In the closed state, Y323 interacts with the main chain of K289ʹ. All distances are presented in Å. **(E)** PLP crater of TA_2799. **(F)** PLP crater of TA_5182 The tyrosine residue fills the crater. The prime mark indicates the residues from another subunit of a dimer.

### Evaluating the role of conserved Y322 in TA_5182 in stabilizing PLP

In the same F321 roof loop, the adjacent Y322 of TA_5182 was also observed for its orientation after PLP binding. The Y322 is initially in H-bond with T127. As the loop gets rearranged, the Y322 moves ∼16 Å to position itself just below the PLP molecule, and the polar OH head group of Y322 interacts with the main chain of the catalytic K289ʹ, stabilizing the catalytic residue (Figure 9D). The orientation of Y322 creates a floor under the PLP molecule, covering the deep crater and allowing better orientation of the PLP molecule to form the internal aldimine. In TA_2799, there is a C324 at the same position. C324 is not adequate to cover the crater, so the crater in the crystal structure of TA_2799 is filled with solvent molecules instead (Figure 9E and F).

Based on this observation, we generated the 2799_L322F_C324Y double mutant. However, this mutant did not show any yellow colour after purification, indicating that the affinity of this double mutant enzyme towards PLP got significantly diminished in comparison to the single mutant 2799_L322F. The enzyme also precipitated within 72h of incubation at 4°C without the addition of cofactor or glycerol. The K_M_ of the double mutant for (*S*)-MBA was 1.188 ± 0.122 mM, which indicates that the affinity of this mutant toward the amino donor substrate has decreased slightly compared to the L322F mutant. The V_max_ and *k*_cat_ values also decreased to 0.033 ± 0.001 U mg^−1^ and 1.381 ± 0.049 min^−1,^ respectively. There was a slight increase in the *k*_cat_/K_M_ (1.162 mM ^−1^ min^−1^) in comparison to the wild-type TA_2799. (Supplementary table 1).

Since this mutation did not increase the affinity of the enzyme towards PLP, we tried mutating the enzyme TA_5182 at position 323, which is mostly a conserved tyrosine or phenylalanine in known bacterial transaminases, to cysteine as found in TA_2799. However, this mutant failed to show any detectable activity against R-PAC, and thus was not studied further for its implication in PLP binding.

### The role of N119 in TA_5182 in PLP binding

Along with the conserved F321 and E123 residues that form an inter subunit network, the presence of the asparagine residue plays a key role in increasing the affinity of the enzyme towards its cofactor. The F321 (L322 in TA_2799) is conserved in many annotated omega-transaminases, but present reports do not describe the formation of yellow colour due to the presence of PLP in the active site of the purified enzyme. TA_5182 has a phenylalanine at the same position, but shows a slight yellow colour at high concentrations above 30 mg/mL. We were interested to understand the reason behind the enhanced affinity of the 2799_L322F mutant towards the cofactor. In the crystal structure of the TA_2799_L322F mutant enzyme, we observed that the ND1 of the N121 residue maintains hydrogen bond with both the OE1 and OE2 of E123, which provides further stability to the anion-pi stacking interaction between E123 and F322 in the same monomer. Apart from that, the OE1 of E123 forms a hydrogen bond with OG of S120, forming an extensive inter-subunit network that facilitates stability of the dimer (Figure 10B). In the crystal structure of the omega-transaminase from *C. violaceum* (PDB ID: 4A6T) and *V. fluvialis* (PDB ID: 4E3R), a similar network is seen. The authors report that the purified protein appears yellow in solution, which supported our observation. On evaluating reported structures of omega-transaminases from various Pseudomonas sp., we found that the ω -TA from *P. jensenni* (PDB ID: 6G4B) harbors an asparagine at this position, and has a Tm_app_ of 62°C, quite similar to the TA_2799_L322F mutant (25). All other reported ω-TAs from other *Pseudomonas* strains, like Pseudomonas str. AAC (PDB ID: 5TI8 and 4UHM) (22, 24), P. aeruginosa (PDB ID: 4B98) (21) and *P. putida* (PDB ID: 6HX9) (17) contains a glycine at this position, like TA_5182. Upon introduction of glutamine at position 119 in TA_5182 replacing glycine, we were able to significantly increase the affinity of the enzyme TA_5182 towards PLP, as observed from the yellow coloration of the purified protein and spectral scan data (Supplementary figure 3B and 4B). In the crystal structure of the ω-TA from *P. fluorescens* (PDB ID: 6S54) (14), there is a leucine and aspartate instead of the asparagine and glutamine as seen in TA_2799. The authors report that the enzyme shows no increase in its affinity towards PLP by the V129N mutation at the PLP binding pocket. A possible reason for such observation can be explained by our study, as the inter subunit network as described in Figure 10A is missing in the TA enzyme from *P. fluoroscens*.

**Figure 10:**
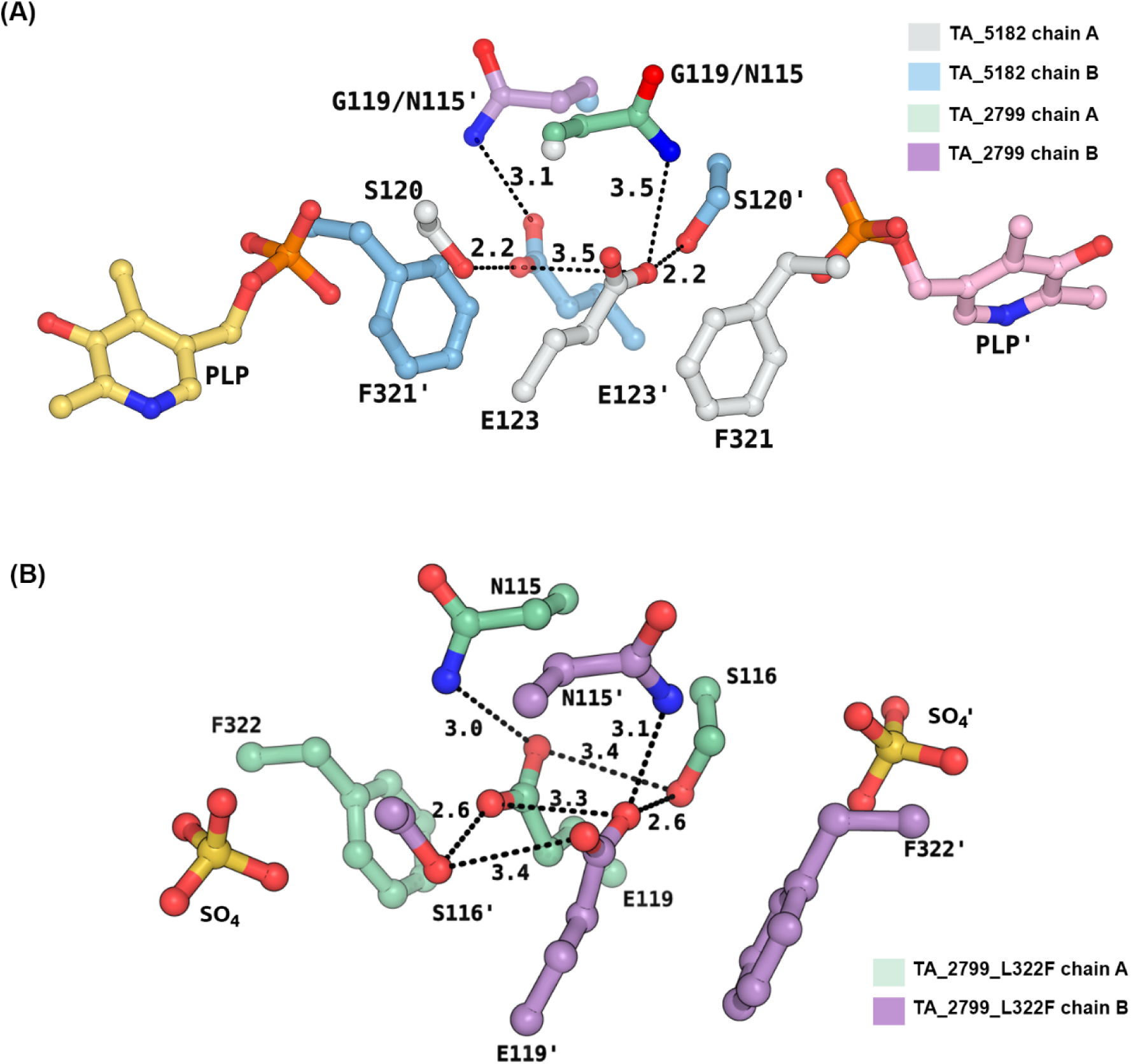
**(A)** Superimposition of TA_2799 and TA_5182 in its closed state showing possible hydrogen bonding network between the sidechains of S120, E123 and N119. The chains A and B of TA_5182 are colored in lightblue and lightgrey respectively. The N115 of TA_2799 from chains A and B are coloured in green and purple respectively. In contrast, TA_5182 contains a glycine at the 119^th^ position. The PLP molecule represents the position of the active site of the enzymes. **(B)** Hydrogen bonding network between the sidechains of S116, E119 and N115 in TA_2799_L322F mutant. The chains A and B are coloured in green and purple respectively. The sulphate ions represent the position of the PGBC of the enzyme. The prime mark indicates the residues from another subunit of a dimer.

## Conclusions

Our study on two ω-TAs from *P. putida* KT2440 provide structural insights into the PLP affinity of bacterial transaminases. We solved a high-resolution cofactor bound structure of TA_2799. We crystallized TA_5182 in two distinct conformations, in which we were able to solve the structure of one of the highly flexible loops that plays a direct role in accommodation of the PLP cofactor in the active site of ω-TAs. This loop is not observed in most of the reported apo forms of the enzyme. The structural information obtained from these crystallographic models helped us investigate the key interactions between the amino acid residues around the cofactor binding pocket of the enzymes. The cumulative effects of these interactions enhance the binding of the PLP cofactor to the enzyme, which in turn stabilizes the overall fold and restricts the loop movement of the enzyme. This is observed by the increase in the melting temperature of the mutants of both the enzymes, along with a significant enhancement in their enzyme activity with *(S)*-MBA and (*R*)-PAC as substrates. The F321 in TA_5182 undoubtedly plays a significant role in accommodation of the phosphate moiety of the cofactor molecule. Apart from this residue, we were able to identify an asparagine residue (N119 in TA_2799) which participates in hydrogen bonding interaction and forms an intricate inter-subunit network that enhances the PLP affinity as well as the dimer stability of the transaminase enzymes. Taken together the structural data, mutations and biochemical studies, the results of this study provide a detailed understanding about the development of stable and efficient ω-TAs for biotransformation.

## Materials and methods

### Lysis and elution buffers

Two types of buffers were prepared for protein purification and subsequent crystallization screening. Lysis Buffer 1: 50 mM Sodium Phosphate (pH 8.0), 300 mM NaCl; Lysis Buffer 2: 50 mM HEPES (pH 8.0), 300 mM NaCl. Corresponding elution buffers for Ni-NTA affinity chromatography contained 250 mM imidazole in addition. TA_5182 was purified in Lysis Buffer 1 for setting up crystallization trays. For all other studies, Lysis Buffer 2 was used.

### Protein expression and purification

The genes coding for *P. aeruginosa* KT2440 ω-TAs TA_2799 and TA_5182 were inserted into pET43.1b-expression vectors, downstream of the 6x-His Tag (Satpute, 2017) and transformed into *E. coli* BL21 DE3 cells by heat shock. Overnight grown *E. coli* cultures were transferred to 5 L baffled Erlenmeyer flasks containing 1.5 L LB (Luria Bertani) broth supplemented with 100 μg/mL Ampicillin and induced with 300 μM Isopropyl thio-β-galactoside (IPTG) at an OD_600_ of 0.9 for overexpression of the proteins. Cells were harvested by centrifugation after 5 h of induction at 24°C in mild shaking conditions (100 rpm) and resuspended in lysis buffer (L1 or L2) and homogenized by sonication. The TA enzymes were purified by affinity chromatography using His Trap Ni-NTA affinity column (GE Healthcare) in the aforementioned buffers. The eluted fractions were concentrated to 15 mg/mL with subsequent buffer exchange with the corresponding lysis buffers to remove imidazole.

### Sequence alignment and conserved residue analysis of TA_2799 and TA_5182

The sequences of TA_2799 and TA_5182 were procured from Uniprot database and aligned using Clustal Omega (30). The conserved regions (highlighted in red) were aligned using the ESpript 3.0 web module (https://espript.ibcp.fr) (31). For analysis of the conserved regions in bacterial Fold type I ω-TAs, sequences of bacterial Fold type I ω-TAs available in the PDB were analyzed with Weblogo tool (https://weblogo.berkeley.edu) (32).

### Site Directed Mutagenesis of TA enzymes

Each PCR reaction mixture consisted of DNA template (pET 43.1b-TA_2799, 60 ng), forward and reverse primer (10 μM, 6.75 μL each), 5x Q5 reaction buffer (10 μL), Q5 GC enhancer (10 μL), Q5 DNA Polymerase (1 μL) and water added to obtain the final reaction volume of 50 μL. PCR was performed as follows: 98℃ for 10 min; 30 cycles of 98℃ for 1 min, 64-72°C (annealing) for 30 s and 72℃ for 10 mins; a final extension at 72℃ for 10 min. 5 μL of 5x FastDigest buffer (Thermo Fisher Scientific^TM^) was added to the PCR product and digested with 1 U of *Dpn*I by incubating at 37℃ for 2 h. The digested product was directly used for transforming competent *E. coli* DH5α bacteria by heat shock for 2 min. The transformed bacteria were spread on LB agar plates supplemented with 100 μg/mL Ampicillin and incubated overnight. Isolated colonies were grown in Ampicillin containing LB media and plasmids were isolated using the Exprep Mini Plasmid Purification Kit (Genexy). The desired mutations were verified by sequencing, and *E. coli* BL21 DE3 cells were transformed using the purified plasmids. The primers used for the SDM studies are listed in Table 3.

**Table 3:**
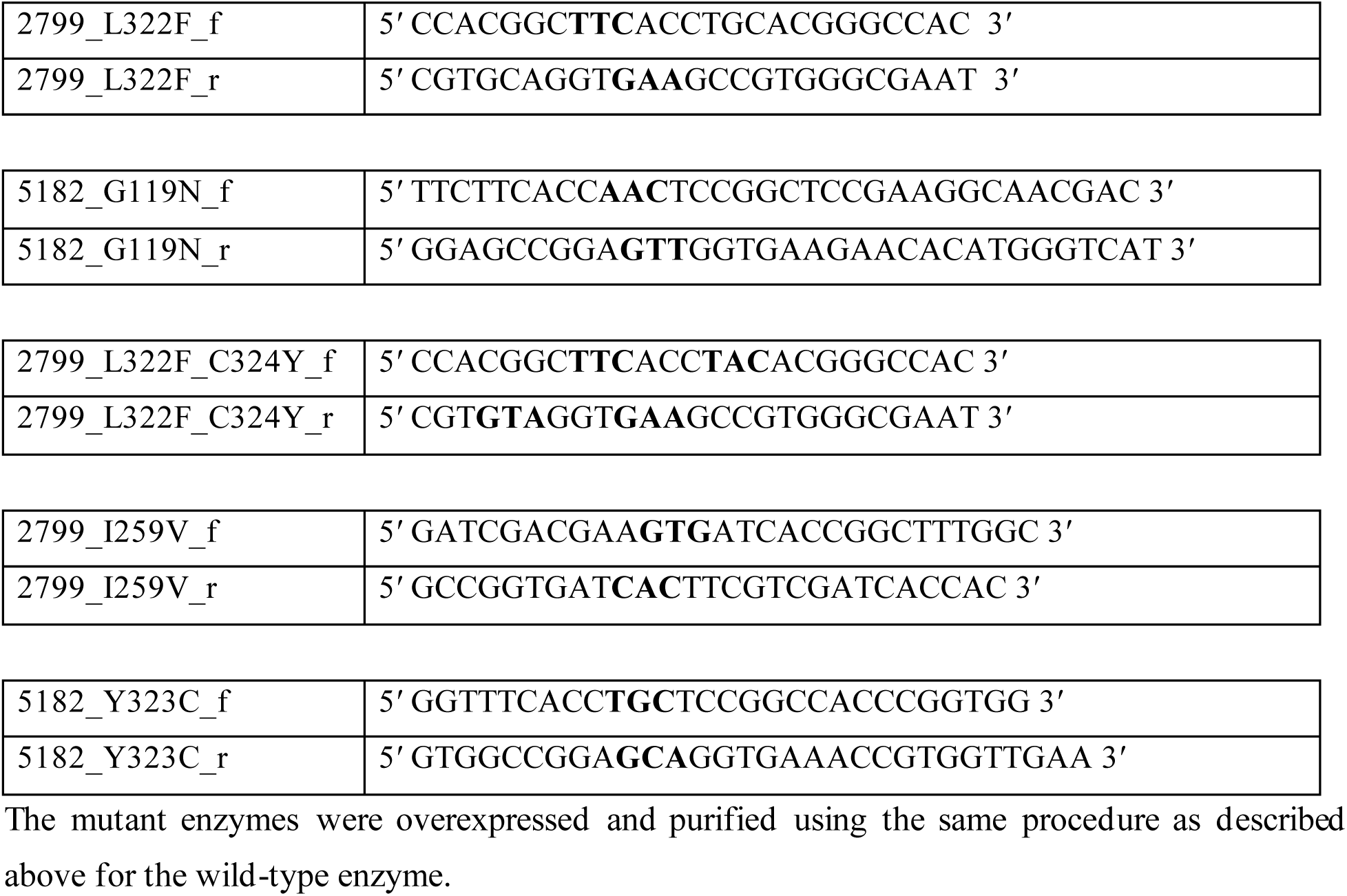
Primers for Site Directed Mutagenesis of the TA enzymes.

### Enzyme activity assay

The triphenyl tetrazolium chloride (TTC)-based assay reported by Sehl et al.(8) was used to investigate the activity of the TA enzymes and their mutants. An 80 µL reaction volume containing 6.25 mM of (*R*)-PAC and 12.5 mM (*S*)-MBA was setup in reaction buffer (50 mM HEPES pH 8.0, 400 μM PLP). To it, 20µl of enzyme (10mg/mL) was added, to make a final reactant concentration of 5 mM amine acceptor ((*R*)-PAC) and 10 mM of amino donor ((*S*)-MBA) in a 100 µL reaction volume. For the standard curve, 100 µL of (*R*)-PAC in reaction buffer at concentration range of 1 mM to 5 mM were setup. All enzyme assays were done in 96-well flat-bottomed microtiter plates (Eppendorf). The plates were incubated in the dark for 2 h at 30°C. After incubation, all the samples were subjected to 10-fold dilution with distilled water and 40 µL of TTC solution (1 mg/mL of TTC salt in a 1:3 v/v solution of 75% ethanol and 1 M NaOH) was added. The reaction of TTC with unused (*R*)-PAC led to formation of a red tetrazolium salt, the absorbance of which was measured at 510 nm using a spectrophotometer (VantaStar^TM^ BMG LabTech). All assays were performed in triplicate.

To evaluate the substrate tolerance of the TA enzymes and their mutants, the activity of the enzymes was measured at varying concentrations of either the amino donor or the amino acceptor keeping other parameters constant. To evaluate the efficiency of the enzymes towards (*R)*-PAC, enzyme activity was measured at varying concentrations of (*R*)-PAC at a fixed concentration of 25 mM (*S*)-MBA and 1 mM PLP. Likewise, the efficiency of the enzymes towards (*S*)-MBA was measured at varying concentrations of (*S*)-MBA at a fixed concentration of 5mM (*R*)-PAC and 1mM PLP.

Temperature dependent conversion of (*R*)-PAC was measured at different temperatures ranging from 25°C to 60°C. The enzymes were incubated with the reaction mixture for 1.5 h followed by denaturation by heating the assay mixture at 90°C for 5 mins and flash cooling in ice. The denatured enzyme was separated from the mixture by centrifuging at 10000g for 3 min, and 10 µL of the supernatant was aliquoted for estimation of remaining (*R*)-PAC.

For pH dependent studies, different buffers corresponding to a pH range from 5.0 to 9.5 were used to prepare the reaction mixtures containing 6.25 mM (*R*)-PAC, 12.5 mM (*S*)-MBA and 500 mM PLP. 20 µL of the purified enzyme (2mg/mL) was added to each reaction and incubated for 1.5 h followed by denaturation by heating the assay mixture at 90°C for 5 mins and flash cooling in ice. The denatured enzyme was separated from the mixture by centrifuging at 10000g for 3 min, and 10 µL of the supernatant was aliquoted for estimation of remaining (*R*)-PAC.

For kinetic studies, the first half of the transamination reaction was measured using High Performance Liquid Chromatography (HPLC). Increasing concentrations of (*S*)-MBA (0-5 mM) was used in the reaction mixture containing 20 mM pyruvate as amino acceptor, 2 mM PLP, and 200 µg of TA enzyme in 100μL reaction buffer. After 30 mins of incubation at 30°C, the reaction was stopped by heating the mixture to 90°C, and 400µL of ddH_2_O was added to the reaction. The mixture was centrifuged at 16000g for 20 min to remove the denatured enzyme. 10µL of the supernatant was injected into the HPLC (Jasco MD-4015) machine at a flow rate of 1mL/min through an Agilent C-18 column. The mobile phase was 30:70 Acetonitrile: ddH_2_O, and the amount of acetophenone produced was measured at 245nm. 1 U of enzyme denotes the amount of enzyme that produces 1 µmol acetophenone per minute.

### Time dependent PLP conversion assay

For measuring the conversion of PLP by the TA enzymes and their mutants, 100 μL reaction were prepared in HEPES buffer pH 8.0, 1 mM PLP and 10 mM (*S*)-MBA. 100µg of enzyme was added to each reaction. The assays were performed in 96-well flat-bottomed microtiter plates (Eppendorf) and readings were taken at every 10 min interval, for two hours. The spectra were recorded from 300 nm to 500 nm.

For PLP dependent conversion of (*R*)-PAC, 100 μL of reactions were set up in flat bottom 96-well plates consisting of 5 mM (*R*)-PAC, 10 mM (*S*)-MBA, 100 μg of enzyme and different concentrations of PLP ranging from 0-1000 μM. The plates were incubated at 30°C for 4 h and the remaining (*R*)-PAC in the reaction mixture was measured using the TTC based method mentioned earlier.

### Thermal shift assay

Fluorescence based assay was used to determine the Tm_app_ values of the TA enzymes and their mutants. 100 μL of the protein solution at a final concentration of 10mg/mL with different solvents were prepared in 50mM HEPES buffer (pH 8.0). 22.5 μL of each of these solutions and 2.5 μL of 50x stock solution of SYPRO orange were added to Genaxy Low Profile PCR (Polymerase Chain Reaction) tubes, mixed thoroughly, and sealed with the provided caps. The assay was performed in BioRad Real Time PCR machine (Bio-Rad, USA) with a linear gradient of increasing temperature (1°C/min) from 15°C to 85°C. The temperature at the maximum rate of fluorescence change (dRFU/d*T*) was taken as the Tm_app_.

### Storage stability assay

Freshly purified TA enzymes and their mutants were concentrated to 10 mg/mL and stored in the dark at 30°C for period of 96 h. Conversion of 5mM (*R*)*-*PAC in the presence of 10 mM (*S*)*-*MBA was measured at periodic intervals in 100 µL reaction assay volume as described above.

### Crystallization

The enzymes TA_2799, TA_5182 and TA_2799_L322F were concentrated to 15-20 mg/mL for setting up crystallization trials. In case of TA_2799, additional glycerol at a final concentration of 12% was added to inhibit precipitation of the enzyme. Crystallization screens were set up using the sitting drop vapour diffusion method with automated Phoenix (Art Robins) robot (Protein Crystallography Facility, IIT Bombay) in 96 well Intelliplate^TM^ with a 1:1 mother liquor: enzyme ratio. The different commercially available screens such as PEG suite (Qiagen), PEG Rx (Qiagen), PEG Ion (Qiagen), Index™ and JCSG+ (Molecular Dimensions) were used for crystallization screening. The crystallization trays were incubated at 22°C (except TA_2799, which was incubated at 18°C) in a cooling incubator. The drops were regularly monitored for crystal growth under a stereomicroscope. Several conditions produced crystals, and the best crystals were grown within 51 days (Supplementary figure 2).

### X-ray diffraction data collection and processing

X-ray diffraction experiments were performed under liquid-nitrogen cryoconditions at 100 K. 25% (v/v) glycerol with the respective mother liquor served as the best cryoprotectant freezing for the crystals. The crystals were briefly soaked in their corresponding cryoprotectant solutions using a nylon loop and flash frozen in a liquid nitrogen stream at 100 K prior to data collection. Data sets of TA_2799 and TA_5182 (open and closed) were collected by the rotation method with 0.5° rotation per frame at a wavelength of 1.5418 Å using Cu *K*α X-ray radiation generated by a Rigaku Micromax 007HF generator equipped with R-Axis IV++ detector at the Protein Crystallography Facility, IIT Bombay. The crystal of TA_2799_L322F mutant was diffracted at the INDUS II synchrotron beamline PX-BL-21 equipped with a MarCCD detector, at RRCAT, Indore at a wavelength of 0.9789 Å. The image frames of the data set were indexed, integrated and scaled using XDS (33). The intensities were converted to structure factors with F2MTZ and CAD in CCP4 (34). The data collection and refinement statistics are summarized in Table 1.

### Structure solution and refinement

The structures of the open and the closed states of TA_5182, and the PLP bound TA_2799 and the TA_2799_L322F mutant were solved by the molecular replacement method. The initial phases were obtained using the MORDA pipeline of the CCP4 online server (35). The model building was done by visual inspection of electron density in COOT (36) and iterative cycles of refinement by REFMAC5 (37). The covalently bound PLP was built in the *F_o_-F_c_* omit electron density map and subsequently refined. The solvent molecules and ions were subsequently added at peaks of electron density higher than 3σ in σ-A weighted *F_o_-F_c_* electron density maps while monitoring the decrease of *R*_free_ and improvement of the overall stereochemistry and Figure of Merit (FOM).

## Supporting information

Supplementary media 1

Supplementary media 2

Supplementary media 3

## Abbreviations

TA: Transaminase
ω-TA: Omega transaminase
PLP: Pyridoxal 5ʹ-phosphate
PMP: Pyridoxamine 5ʹ-phosphate
PGBC: Phosphate group binding cup
(*R*)-PAC: (*R*)-Phenylacetylcarbinol
RMSD: Root mean square deviation
(*S*)-MBA: (*S*)-Methylbenzylamine
TTC: Triphenyl tetrazolium chloride

## Data availability

The atomic coordinates of structures of TA_2799, TA_2799_L322F, TA_5182 open and TA_5182 closed were deposited in the Protein Data Bank (PDB) with accession codes 9J2K, 9J4Y, 9J4Z and 9J50 respectively. All relevant data are available from the corresponding author upon request.

## Author contributions

P.B., S.B.N. and P.D. conceptualization, visualization and analysis. P.D. investigation. P.D. and P.B. writing: original draft. P.B. and S.B.N. supervision and writing: review and editing.

## Acknowledgements

We thank Dr. Ravindra Makde, Dr. Biplab Ghosh and Dr. Ashwani Kumar for their support in diffraction data collection at the PX-BL-21 beamline in INDUS II synchrotron at Raja Ramanna Centre for Advanced Technology (RRCAT), Indore. We acknowledge the “Protein Crystallography Facility” at IIT Bombay for the crystallization and X-ray diffraction data collection. P.D also thanks the Council of Scientific & Industrial Research (CSIR), Ministry of Science and Technology, Govt. of India for providing the Ph.D. fellowship (file no. 09/087(0975)/2019-EMR-I) for the research.

