## Supplementary Information for "Structural insights and rational design of *Pseudomonas putida* KT2440 Omega transaminases for enhanced biotransformation of (*R*)-Phenylacetylcarbinol to (1*R*, 2*S*)-Norephedrine"

**Running title:** Rational design of omega transaminases

[*]To whom correspondence should be addressed:

Santosh Noronha, Department of Chemical Engineering, Indian Institute of Technology Bombay, Powai, Mumbai-400076, India;

Prasenjit Bhaumik, Department of Biosciences and Bioengineering, Indian Institute of Technology Bombay, Powai, Mumbai-400076, India;

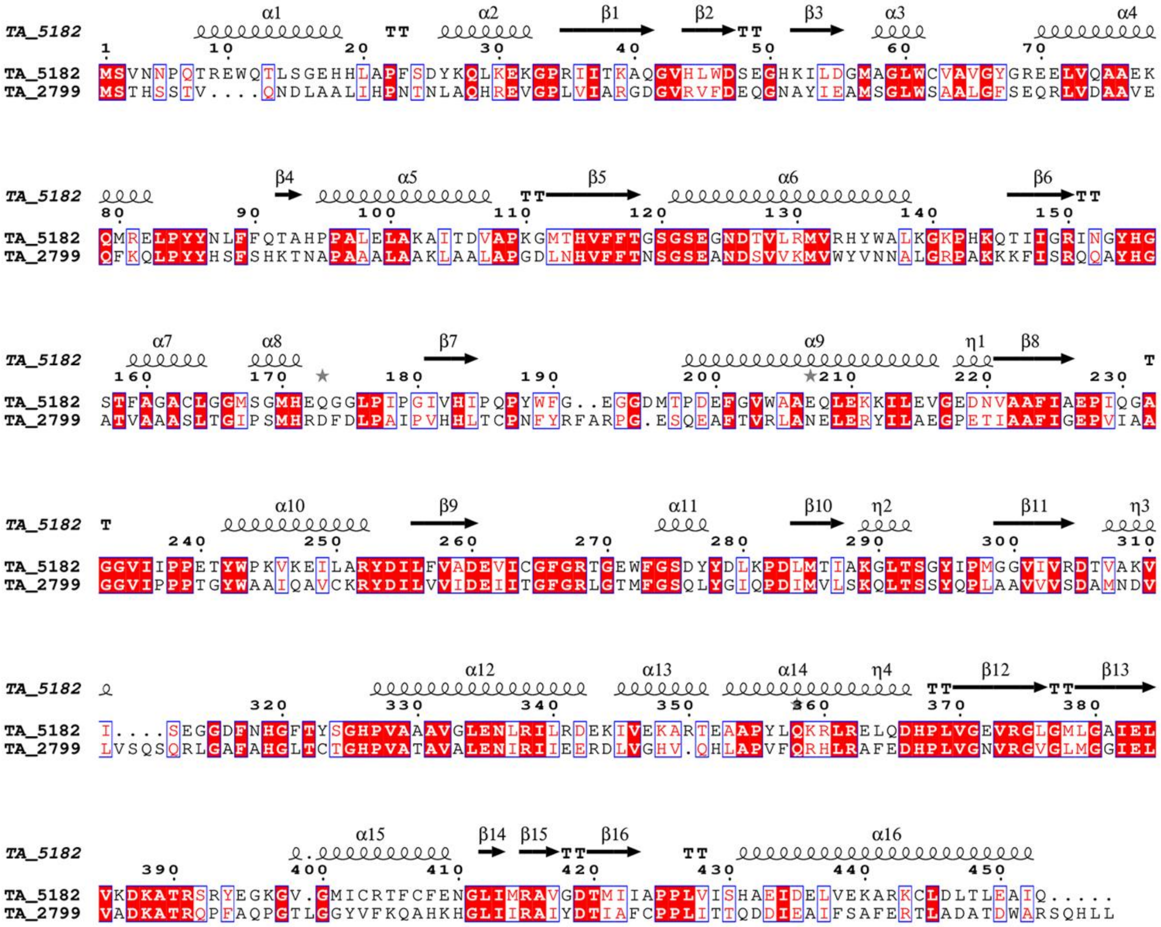

**Supplementary figure 1:** Sequence alignment of TA_2799 and TA_5182. Secondary structural elements of TA_5182 crystal structure is also shown on the top of the alignment.

### Crystallization of the enzymes:

Crystallization screening drops were set in 480 different conditions in buffers with and without PLP. Both the transaminases formed crystals after at least 50 days of incubation. For TA_2799, very tiny crystals were formed in the absence of the cofactor molecule, which did not diffract. On co-crystallization of TA_2799 with 1mM PLP, a single large and robust crystal was formed in the condition 0.2 M Ammonium acetate, 0.1 M Bis-Tris 5.5, 25% w/v PEG 3,350 (INDEX G6), with a visible yellow colour signifying the crystallization of the holo enzyme (Supplementary figure 1). The crystal diffracted to a resolution of 1.67 Å at the corner of the square detector, however, the data was limited to the detector edge at 1.76 Å.

The mutant enzyme TA_2799_L322F was also crystallized in a condition containing 0.2 M Lithium sulphate, 0.1 M bis-Tris pH 5.5, 25 % PEG 3350 (JCSG+ H9). Crystals were observed after 52 days of incubation at 18°C and the crystals diffracted to a resolution of 2.67 Å.

We were not able to crystallize the wild-type TA_5182 in its PLP bound holo form. Co-crystallization with 1 mM PLP did not lead to any crystal formation in the drops. However, TA_5182 was crystallized in two different apo forms. The open state of the enzyme crystallized in the condition containing 0.1 M Bis-Tris pH 5.5, 2 M ammonium sulphate, (INDEX A3 and JCSG+ G11) and these crystals diffracted to a resolution of 3 Å, but cutoff at 3.4 Å. We were also able to crystallize TA_5182 in its closed state, in the condition 1.4 M Sodium Potassium Phosphate, pH 8.2, (Index B7). The data collection and refinement statistics are reported in table 1.

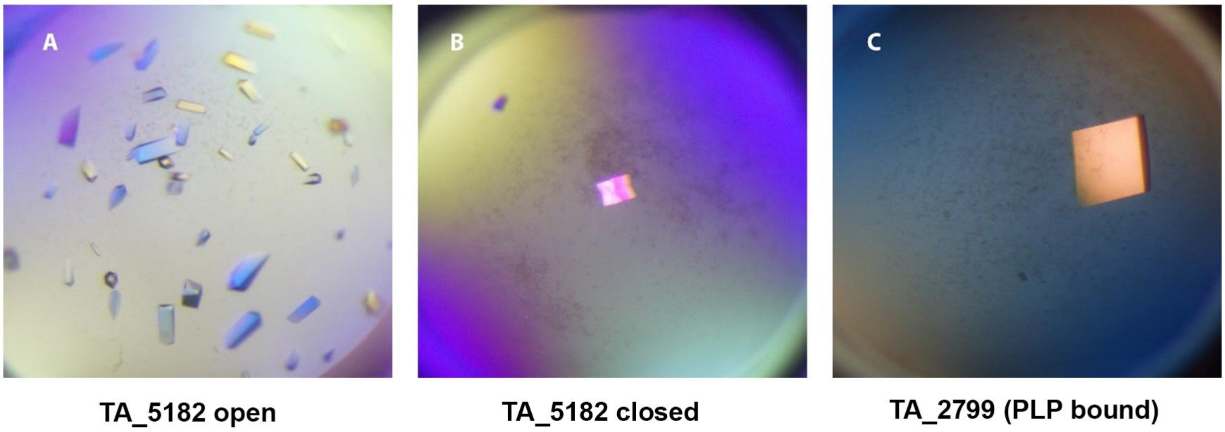


**Supplementary figure 2:** Crystallization drops of **(A)** TA_5182 in open state **(B)** TA_5182 in closed state and **(C)** TA_2799 in PLP bound holo state.

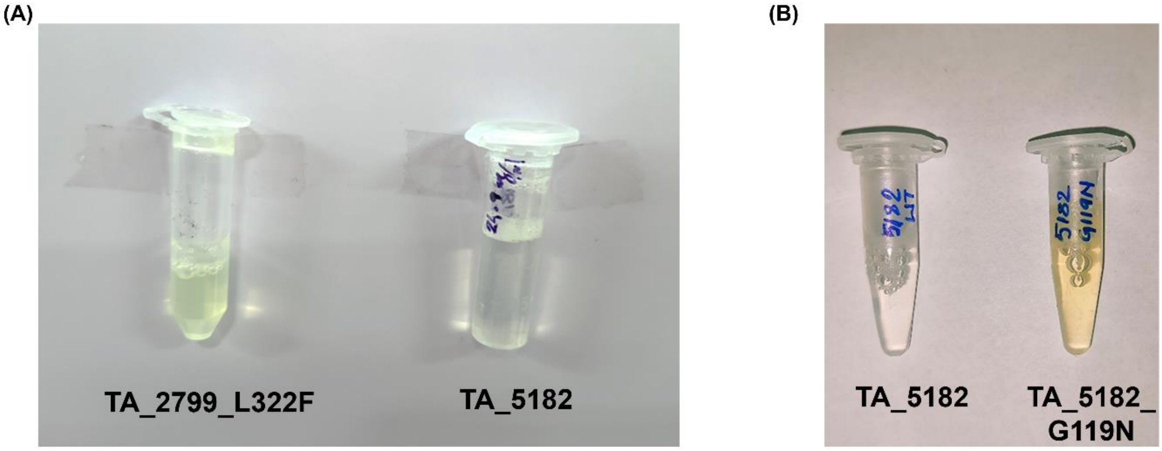


**Supplementary figure 3:** Purified protein showing yellow colouration in **(A)** TA_2799_L322F and **(B)** TA_5182_G119N in comparison with TA_5182.

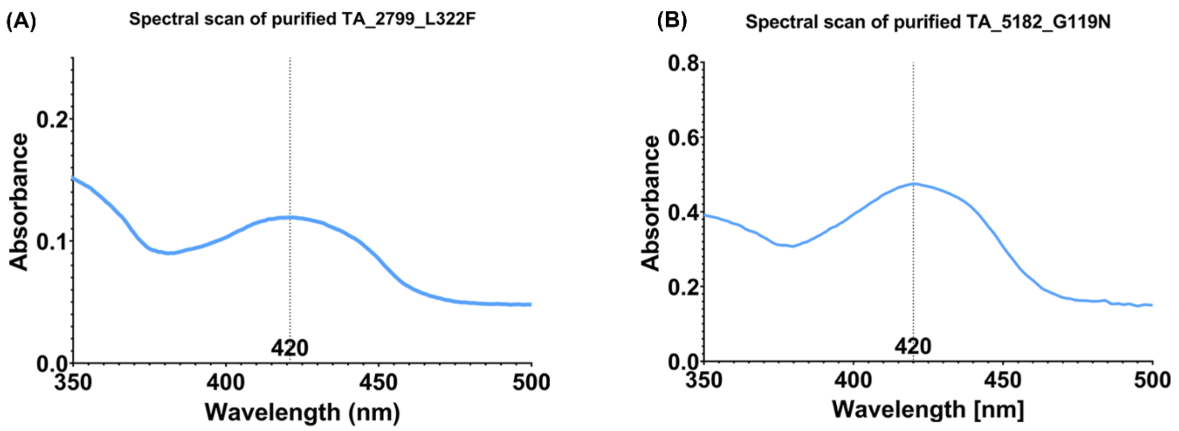


**Supplementary figure 4:** Spectral scan of **(A)** TA_2799_L22F and **(B)** TA_5182_G119N showing absorption peak at 420 nm corresponding to PLP bound form of the protein.

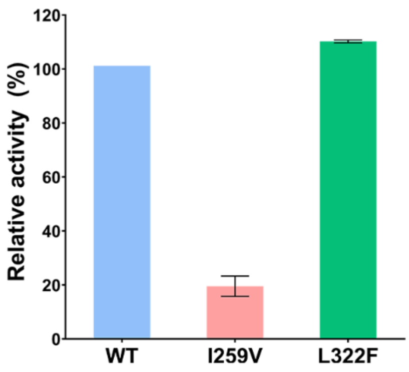


**Supplementary figure 5:** Relative activity of TA_2799 wild type enzyme and its mutants towards conversion of 5mM *(R)*-PAC in the presence of 10mM *(S)*-MBA and 400 µM PLP.

**Supplementary media 1:** Overall structure of PLP bound TA_2799 showing different structural features of Fold type I ω-TAs: i. Cartoon representation of the monomer with covalently bound PLP at the active site. ii. Cartoon representation of the dimer showing the PLP binding pocket and the flexible loops. iii. Surface representation of the dimer showing the position of the flexible loops and the substrate entry channel of the enzyme.

**Supplementary media 2:** Transition of F321 from the open to closed state in TA_5182 leading to conformational changes in the side chains of residues around the PLP binding pocket of the enzyme.

**Supplementary media 3:** Transition of Y323 from the open to closed state in TA_5182 forming hydrogen bond interactions with catalytic K289 in the closed conformation of the enzyme.

**Supplementary Table 1: Kinetic parameters of the TA_L322F_C324Y**

| Enzyme | $K_M$ (mM) | $V_{max}$ (U $min^{-1}$) | $k_{cat}$ ($min^{-1}$) | $k_{cat}/K_M$ ($mM^{-1}$ $min^{-1}$) |
|---|---|---|---|---|
| **TA_2799_L322F_C324Y** | 1.188 ± 0.122 | 0.033 ± 0.001 | 1.381 ± 0.049 | 1.162 |
