## Supplementary figures and images for "Structural insights and rational design of *Pseudomonas putida* KT2440 Omega transaminases for enhanced biotransformation of (*R*)-Phenylacetylcarbinol to (1*R*, 2*S*)-Norephedrine"

### Supplementary media 2

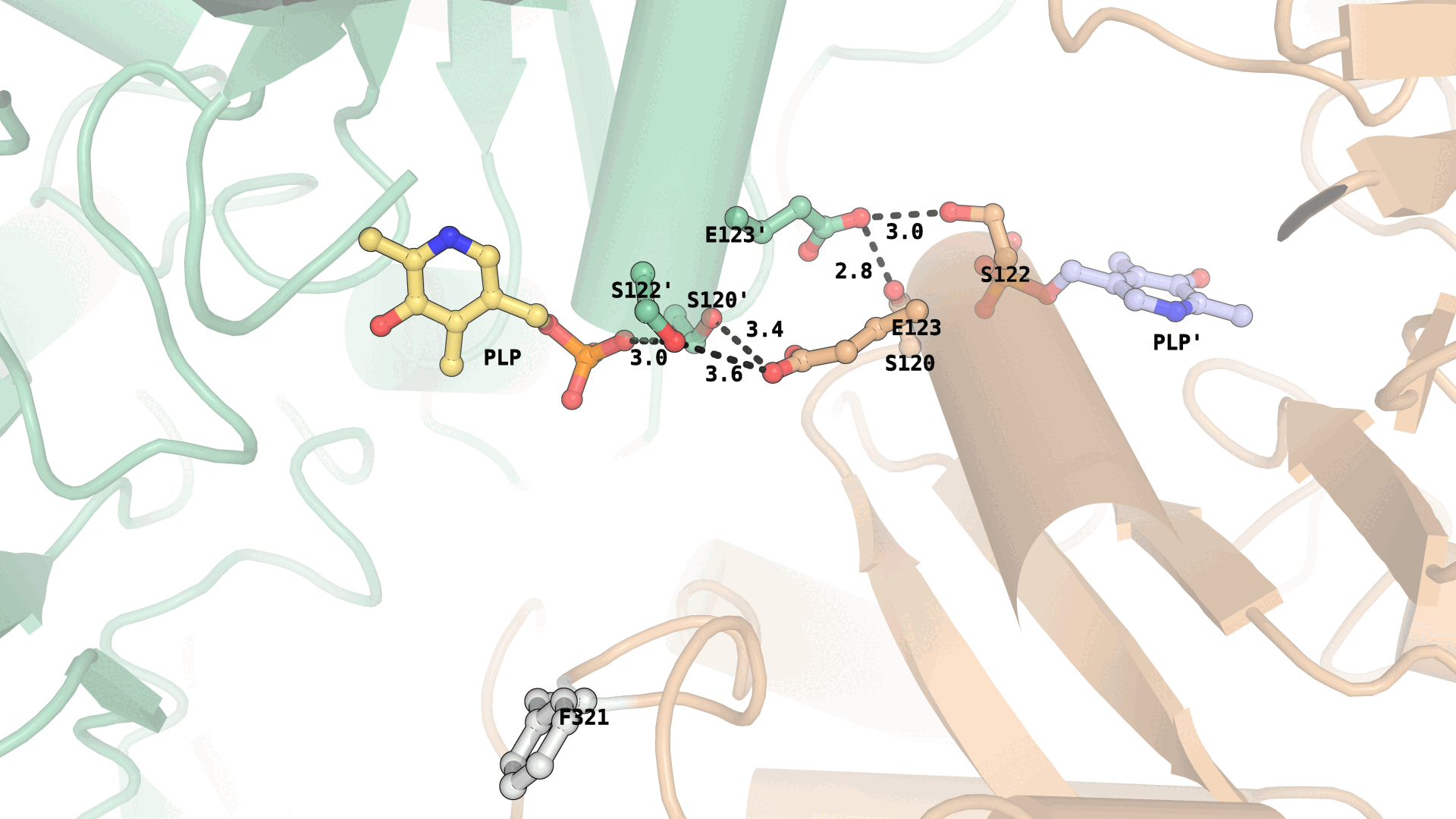

### Supplementary media 3

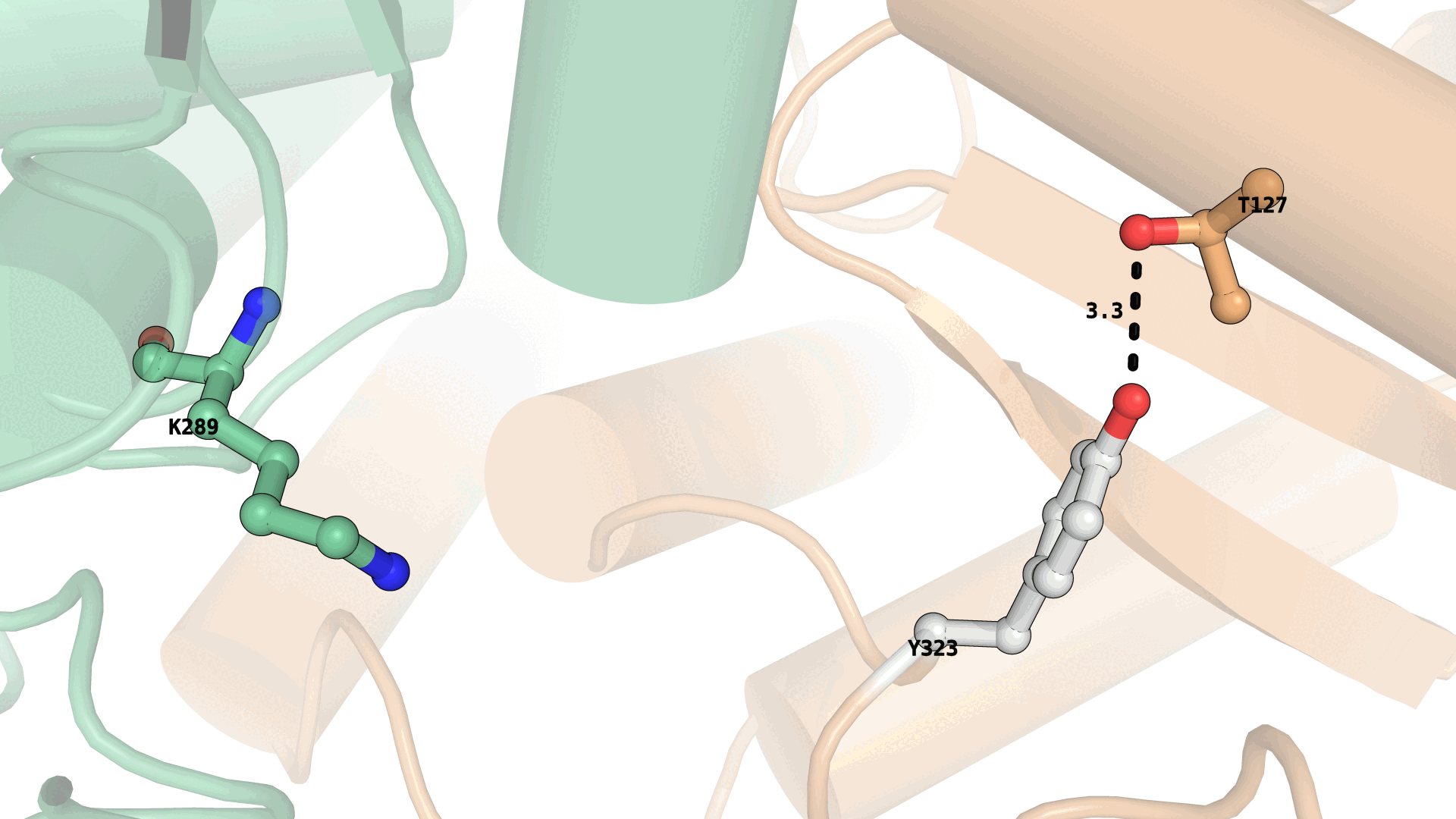
